# Insulation of gene expression by CTCF and cohesin-based subTAD (ubiquitous intra-TAD) loop structures in mouse liver

**DOI:** 10.1101/271023

**Authors:** Bryan J. Matthews, David J. Waxman

## Abstract

CTCF and cohesin are key drivers of 3D-nuclear organization, anchoring the megabase-scale Topologically Associating Domains (TADs) that segment the genome. Here, we present a computational method to identify cohesin-and-CTCF binding sites that form intra-TAD DNA loops (subTADs). We show that predicted subTAD anchors are structurally indistinguishable from those of TADs regarding their binding partners, sequence conservation, and resistance to cohesin knockdown; further, the subTAD loops retain key functional features of TADs, including insulation of chromatin contacts, blockage of repressive histone mark spread, and ubiquity across tissues. We propose that subTADs form by the same loop extrusion mechanism as larger loops, and that their shorter length enables finer regulatory control over gene expression. 4C-seq analysis using an Alb promoter viewpoint illustrates the role of subTADs in restricting enhancer-promoter interactions. These findings elucidate the role of intra-TAD cohesin-and-CTCF binding in nuclear organization, and demonstrate that distal enhancer insulation by subTADs is widespread.

## INTRODUCTION

The mammalian genome is organized into stereotypical domains, averaging ~700 kb in length, called Topologically Associating Domains (TADs) [1, 2]. TADs are insulated chromatin domains whose genomic boundaries are largely retained across tissues [1] and have been conserved during mammalian evolution [3, 4]. TADs provide a stable genomic architecture that constrains enhancer-promoter contacts, while allowing for dynamic tissue-specific interactions that stimulate gene expression within TADs, thereby linking chromatin structure and positioning to gene expression [5, 6].

Hi-C, an unbiased genome-wide chromosome conformation capture method [7], identifies TADs based on their insulation from inter-domain interactions and by the increased frequency of intra-domain interactions that occurs within individual TADs [1, 2]. TADs show substantial overlap with features of nuclear organization identified using other approaches, including replication domains, lamina-associated domains, and A/B chromatin compartments [4, 8, 9]. TADs impact gene expression by insulation, which limits a given gene’s access to regulatory regions [10]. While overall TAD structure is generally shared across tissues within a species, individual TADs often show major tissue-specific differences in their spatial positioning within the nucleus, and in their overall activity, transcription factor (TF) binding patterns, and patterns of expression of individual genes [4]. It is unclear to what extent these large megabase-scale chromatin structures exert regulatory control over the multiple, often variably-expressed, genes found within their boundaries.

Two key protein factors, CCCTC-binding factor (CTCF) and the multi-subunit cohesin complex, underpin nuclear organization at a kilobase scale by cooperatively engaging genomic DNA via a loop extrusion complex, which is dynamically mobile within TAD boundaries and may help organize their structure [11–13]. CTCF is an 11 zinc finger protein that stably binds DNA and can serve as an insulating enhancer-blocker and a modulator of 3D chromatin structure [14]. Sites bound by both cohesin and CTCF (cohesin-and-CTCF (CAC) sites) are associated with insulator function [5, 15] and are found at TAD boundaries [1, 2], while sites bound by cohesin but not CTCF (cohesin-non-CTCF (CNC) sites) are found at tissue-specific promoters and enhancers [16] and may help to stabilize large TF complexes [17]. In addition, topoisomerase-IIβ (Top2b) is associated with CAC complexes at TAD boundaries in mouse liver, presumably to relieve the torsional strain of the extrusion complex [18]. Several lines of evidence indicate that CTCF and cohesin are the primary architects of nuclear organization in mammals [13, 19, 20]. Complete knockout of either CTCF or cohesin is embryonic lethal [21–23], whereas partial depletion of CTCF or cohesin results in altered gene expression but limited phenotypic impact [19, 23, 24]. More recent studies using inducible degradation systems indicate that complete removal of CTCF or cohesin-related factors leads to a loss of loop structures at multiple scales in a highly dosage-dependent manner [25–27]. Mutations affecting CAC loop anchors are frequently seen in cancer and lead to dysregulation of adjacent genes, evidencing the functionality of these loops [28–30]. However, there are many more CAC sites within TADs than at TAD boundaries, and it is not clear what factors differentiate loop-forming CAC sites at TAD boundaries from other CAC sites in the genome.

One of the challenges in connecting Hi-C interaction maps to specific CAC sites is the comparatively low resolution of Hi-C datasets, typically on the order of 25-100 kb. Consequently, Hi-C analysis has primarily been directed to the study of nuclear organization at the level of megabase-scale TAD structures. However, high resolution Hi-C datasets obtained using extreme deep sequencing (>25 billion reads) have led to two key discoveries [12]. First, ~90% of DNA loops (“loop domains”, defined as local peaks in the Hi-C contact matrix) are associated with both CTCF binding and cohesin binding, and 92% of such loops involve inwardly oriented CTCF anchors [12]. Thus, loop anchors are bound at asymmetric CTCF motifs that face the loop interior. This previously unappreciated feature of CTCF loops facilitates the identification of such loops *in silico* [13, 31]. Furthermore, expression of neighboring genes changes in a predictable manner when CTCF anchors are inverted or deleted by CRISPR/Cas9 genomic editing [5, 13, 20]. Second, extreme deep sequencing Hi-C studies identify a much larger number of shorter loops than previously recognized (~10,000 loops with a median size of 185 kb) [12], many of which represent complex nested structures (e.g., isolated cliques) [13]. The ability to distinguish between such substructures has led to predictions ranging from 10^3^ to 10^6^ loops per genome, depending on the 3C-based analysis method and the cutoff values employed [13, 32–34]. The presence of nested loop structures may be a general feature of topological nuclear organization, and the ability to detect such structures is dependent on the method, resolution, and computational approach [32–36].

While sub topologies within TADs have been observed, it is unknown whether those interactions represent enhancer-promoter loops or other looped structures, and whether they are mediated by cohesin, mediator, or other architectural proteins [15, 37]. Short, < 200 kb CTCF-anchored loops, termed chromatin contact domains or super-enhancer domains, have been identified in mouse embryonic stem cells (mESCs) by ChIA-PET experiments that select for CTCF and cohesin binding sites [32, 38], and are enriched for tissue-specific genes and enhancers [5, 32]. However, these genomic regions represent a minority of CTCF-anchored DNA loops, and likely do not fully represent all of the nuclear topological domains evident in high resolution Hi-C maps [12, 26, 39]. Given the inability to identify CAC-anchored intra-TAD loops (subTAD loops) from standard, low resolution Hi-C data, we sought to build on the above advances and develop a computational method to predict and then experimentally validate such subTAD-scale loops by using only 2D (CTCF and cohesin ChIP-seq binding activity) and 1D (CTCF motif orientation) information. Here we define subTADs as intra-TAD loops anchored by cohesin and CTCF that contain at least one gene; these subTADs are mechanistically distinct from short range enhancer-promoter loops, and from longer range genomic compartmentalization [26, 27, 40] whose impact on gene expression is also discussed.

Here we present a computational method to identify subTAD loops genome-wide. We elucidate the structural and functional features of the subTADs identified, and those of the better-established TADs, including their impact on gene expression in a mouse liver model. We show that, mechanistically, subTADs are anchored by loop extrusion CAC complexes that are shared across tissues and show strong conservation. Further, we demonstrate that, at a functional level, subTAD loops insulate repressive chromatin mark spread and thereby enable selective, high-level expression of genes relative to their immediate genomic neighbors, including gene targets of super-enhancers, as well as genes otherwise found in repressive nuclear compartments. These findings reveal how subTADs harness many of the same mechanisms as TADs but in ways that allow for greater local control of gene expression.

## RESULTS

### I. Features of TADs and their functional impact on gene expression

#### Features associated with TAD boundaries

We characterized TADs identified in mouse liver [3] using matched ChIP-seq datasets for CTCF and the cohesin subunit Rad21, which we obtained for a series of individual adult male mouse livers. Genomic regions co-bound by cohesin and CTCF (CAC sites) were strongly enriched at TAD boundaries (Figure 1A), consistent with [3]. In contrast, cohesin-non-CTCF sites (CNC sites) were weakly depleted at TAD boundaries. We also observed strong enrichment for motif-oriented CTCF binding at TAD boundaries (Figure S1A), consistent with recent reports and the loop extrusion model of domain formation [3, 12, 13]. Next, we explored the impact of cohesin depletion on CAC sites associated with TAD boundaries, following up on the finding that many cohesin binding sites are maintained upon knockout or knockdown of components of the cohesin complex [17]. Figure 1B shows the distribution of cohesin-bound regions that are either resistant or sensitive to haploinsufficiency of the cohesin subunit Rad21 in hepatocytes, or to knockout of the cohesin subunit Stag1 in mouse embryonic fibroblasts (MEFs). In both cell types, sites resistant to cohesin loss are enriched at TAD boundaries, while those sensitive to cohesin loss are more equally distributed along the TAD length. This may explain the unexpected finding that domains and compartments are largely maintained after depletion of cohesin [15, 37, 41].

**Figure 1:**
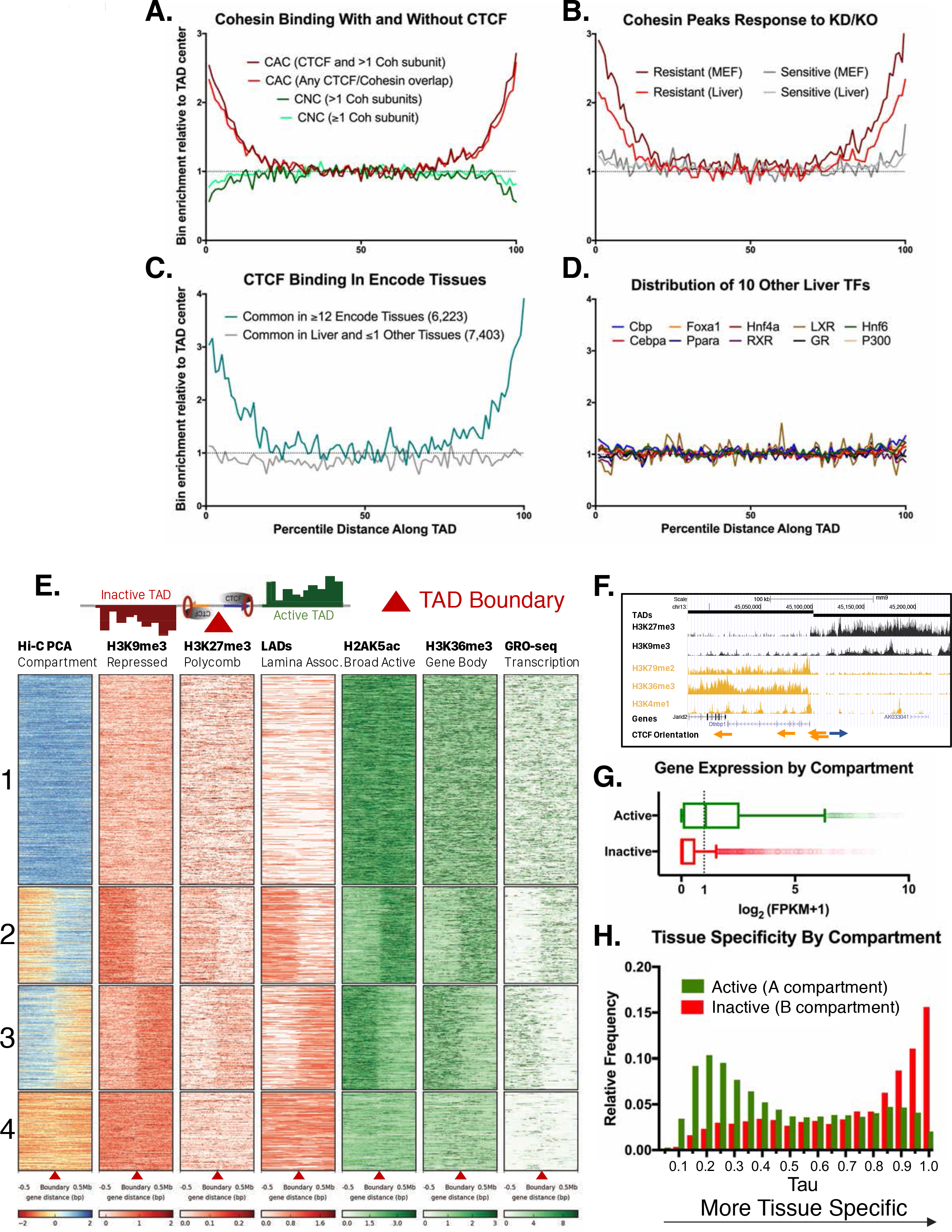
Features of TAD boundaries and TAD insulator function. A. Cohesin-and-CTCF (CAC) sites are enriched at TAD boundaries, while cohesin-non-CTCF (CNC) sites are weakly depleted. As the cohesin (Coh) complex is a multi-protein complex, the darker color within each group represents a stricter overlap of three cohesin subunits (Rad21, Stag1 and Stag2). For Figures 1A-D and Figure S1B-F, profiles indicate a normalized aggregate count of peaks or features along the length of all TADs, sub-divided into 100 equally-sized bins per TAD where bin #1 is the 5’ start of the TAD. Normalization was performed to allow comparison of multiple groups with variable peak numbers in a single figure. As described in the methods, the y-axis indicates the enrichment within a given bin versus the average of the 5 center bins (bins #48-52). B. In both mouse liver and MEFs, cohesin binding sites that are resistant to knockdown (KD) or knockout (KO) of component subunits (~40% of sites) are strongly enriched for TAD boundaries. Cohesin sites that are sensitive to loss following KD or KO (~60% of sites) are not enriched at TAD boundaries. C. CTCF sites that are deeply-shared across ENCODE tissues (≥12 other tissues) are strongly enriched at TAD boundaries, while those that are either unique to liver or shared in one other tissue are not enriched at TAD boundaries. The number of CTCF sites is shown in parenthesis. D. 10 liver-expressed TFs show no appreciable enrichment at TAD boundaries. Of all the publicly available ChIP-seq data tested, these data are representative of the vast majority of the >50 publically available ChIP peak lists for liver-expressed TFs. Notable exceptions related to promoter-associated features, marks, and factors are shown in Figure S1B,D. This enrichment does not appear to be an artifact of CTCF overlap (Figure S1E). E. A subset of TAD boundaries represent transitions from active to inactive chromatin compartments. Shown is the distribution of activating and repressive marks or features for a 1 Mb window around each TAD boundary. TAD clusters, numbered at the left, are defined using k means clustering (k=4). The boundaries between TADs transition from active to inactive genomic compartments (or vice versa) for TAD clusters 2 and 3. For downstream analyses, a TAD was considered active if the boundary at the start of a TAD fell into clusters 1 or 2 (the first two clusters from the top) and the boundary at the end of the same TAD fell into clusters 1 or 3 (see Methods). See Table S1A for a full listing of the 3,538 TADs analyzed and their active/inactive status. F. UCSC browser screenshot for a transitional TAD boundary on chromosome 13 from TAD cluster 3 in Figure 1E. Arrows at bottom indicate CTCF motif orientation. G. Genes found in active compartment TADs are more highly expressed, with the majority of genes showing >1 FPKM, than genes found in inactive TAD compartments. Box plots are based on 12,258 genes in 1,930 active TADs and 4,643 genes in 1,000 inactive TADs. 939 genes in 473 of the inactive TADs are expressed at > 1 FPKM (see Table S1E). Genes in weakly active and weakly inactive TADs were not included in these analyses. H.Genes whose TSS are located in inactive TADs (B compartments) are more tissue specific in their expression pattern than genes found in active TADs (A compartments). The top GO category for expressed genes in the A compartment is RNA binding, while the top category for expressed genes in the B compartment is monooxygenase activity (not shown).

Given the general conservation of TAD boundaries between tissues in both mouse and human [1, 2, 4], we compared regions bound by CTCF in mouse liver to 15 other mouse tissues from the ENCODE Project [42]. CTCF sites that are shared across 12 or more tissues showed 3-4 fold enrichment at TAD boundaries relative to the center of the TAD, whereas CTCF binding sites unique to liver, or shared with only one other tissue, showed no such enrichment (Figure 1C). The TAD boundary enrichment seen for CTCF and cohesin was absent for >50 other liver-expressed TFs whose binding site distribution within TADs we examined (Figure 1D). Tbp and E2f4, which are characterized by promoter-centric binding [43, 44], are two notable exceptions (Figure S1B). Consistent with this, TAD boundaries were enriched for promoters of protein-coding genes, including promoters that do not overlap CAC sites, and for histone marks associated with promoters but not enhancers (Figure S1C-E). TAD boundaries were also enriched for CpG hypomethylation, which was most pronounced at TAD anchor CTCF motifs (Figure S1F-H). CpG methylation is greater at CAC sites not involved in TAD or subTAD anchors (see below), and could represent an additional layer of epigenetic regulation of CAC-based loop formation.

#### TADs segregate the genome into compartmentalized units

TADs have the ability to insulate the spread of repressive histone marks and also enhancer-promoter interactions, referred to as enhancer blocking [1, 5, 37]. By these dual mechanisms, TADs can exert control over tissue-specific gene expression, despite the TADs themselves being largely structurally invariant across tissues. As TADs are defined based on their insulation of chromatin contacts, we investigated their impact on chromatin mark spread. We examined four broad histone marks associated with either repression (H3K9me3, H3K27me3) or activation (H2AK5ac, H3K36me3). We also examined Global Run-on Sequencing data to identify actively transcribed regions of the genome, as well as Lamina Associated Domain (LAD) coordinates to visualize areas of the genome associated with the nuclear periphery. Figure 1E shows a heat map representation of a 1 Mb window around each TAD boundary, clustered using k-means clustering (k=4) based on H3K9me3 and H2AK5ac ChIP-seq data and on the Eigen value of the Hi-C prinicipal component analysis (PCA), which provides an estimate of active versus inactive genomic compartments [7]. A subset comprised of 1,439 TAD boundaries (40.9% of all boundaries) represents transitions from inactive to active chromatin compartments, or vice versa (Figure 1E; 2^nd^ and 3^rd^ clusters). Also shown is an example of a transitional TAD boundary on chromosome 13, with a shift from active to inactive chromatin marks separated by inversely-oriented CTCF binding sites (Figure 1F). Using the clusters shown in Figure 1E, each TAD was designated as active, weakly active, inactive, or weakly inactive, based on the signal distribution around the boundary and the eigenvalue from Hi-C PCA analysis along the length of the TAD (see Suppl. Methods and listing of TADs in Table S1A). Striking differences in gene expression were seen between active and inactive compartment TADs (median expression 1.095 FPKM for 12,258 genes in active TADs vs. 0.003 FPKM for 4,643 genes in inactive TADs; Figure 1G).

We sought to determine the tissue-specificity of the genes in active vs. inactive TADs. We used the expression level of each gene across ENCODE tissues to calculate Tau scores, a robust metric for tissue specificity [45, 46]. Tau values are close to 1 are highly tissue specific, while lower values (< ~0.3) are widely expressed and considered housekeeping genes (see Methods). A greater fraction of genes located in inactive TADs are tissue-specific compared to genes in active TADs (Figure 1H). Overall, only 939 (20.2%) of all 4,643 genes in inactive TADs are expressed in liver (FPKM >1) vs. 6,290 (51.3%) of the 12,258 genes in active TADs. Furthermore, genes whose TSS is close to a TAD boundary (i.e., TSS within 2% of the total TAD length in either direction from the boundary) tend to be less tissue specific than the genomic average (Figure S1H). Active transcription may be a key driver of dynamic cohesin movement in the nucleus [47], and RNA polymerase II, *in vitro*, is capable of translocating cohesin rings along DNA [48]. Thus, the ubiquitous expression of genes at TAD boundaries could be either a driver or an initiator of loop extrusion, although the exact mechanism remains unknown.

### II. Identification of subTAD loops

#### Predicting subTAD loops from TAD-internal CTCF and cohesin binding sites

While CAC sites that are tissue ubiquitous, cohesin knockdown-resistant, or species-conserved show a clear 2 to 5-fold enrichment at TAD boundaries (Figure 1), a large majority of such sites are TAD-internal and presumably do not contribute to TAD formation. Overall, only 14.7% of liver CTCF binding sites are associated with TAD boundaries (Figure S2A), consistent with other reports [1], and only 23% of the CTCF-bound regions that retain all four of the above features are within 25 kb of a TAD boundary (Figure S2A). We considered two possibilities: 1) TAD-internal CAC sites form subTAD loops that are too short to be detected in standard Hi-C datasets; and 2) additional factors associated with TAD boundary CTCF sites differentiate them from other such binding sites in the genome (see Discussion). To examine the first possibility, we used an algorithm for analysis of CTCF loops [31] and adapted it to predict subTAD-scale loops *in silico*, using CTCF and cohesin peak strength and CTCF orientation as inputs (Figure 2A), and building on the finding that >90% of CTCF-based loops are formed between inwardly-oriented CTCF sites [12]. Each mouse liver CAC site was given a score that represents its peak strength and CTCF motif score, and an orientation was assigned based on whether the non-palindromic CTCF motif was present on the (+) strand or the (−) strand, considering the highest scoring CTCF motif at each CAC site. Scanning the genome, each (+) strand CAC peak was connected to putative downstream (−) strand CAC sites. Low scoring CAC peaks were removed and the process was iteratively repeated until the top 20,000 candidate subTAD loops remained. The set of loops was then filtered, as detailed in Methods, to take into account cohesin scoring, and to ensure TSS overlap and <80% TAD overlap, to restrict our definition of subTADs to intra-TAD CAC-mediated loops that contain at least one TSS. Applying this approach to each of 4 matched pairs of liver ChIP datasets for CTCF and cohesin, we identified a set of 9,543 subTAD loops present in all four liver samples, with a median subTAD length of 151 kb. These subTADs differ substantially from the generally shorter and much larger number of CTCF loops predicted by the original algorithm [31] (Figure S2B; see Methods). Functionally, anchors of these shorter loops show less insulation and weaker directional interactions than the subTAD anchors identified in our study (Figure S2C,D; also see below). Moreover, 91% of our predicted subTAD loops were wholly contained within a single TAD, versus only 67% for a random shuffled control (Figure S3A). Figure 2B illustrates subTAD structures within TADs along a segment of chromosome 2 and highlights their substantial overlap with “CTCF-CTCF” DNA loops identified by ChIA-PET analysis of cohesin-mediated interactions in mouse embryonic stem cells (mESCs) [5]. The final set of 9,543 subTADs includes 1,632 subTADs (17.1%) that share one CTCF loop anchor with a TAD boundary (i.e., subTAD nested in a TAD with a potential shared anchor). Within TADs we observe single or nested subTADs, as well as greater subdivision of larger TADs into multiple subTAD structures (Figure S2E-G).

**Figure 2:**
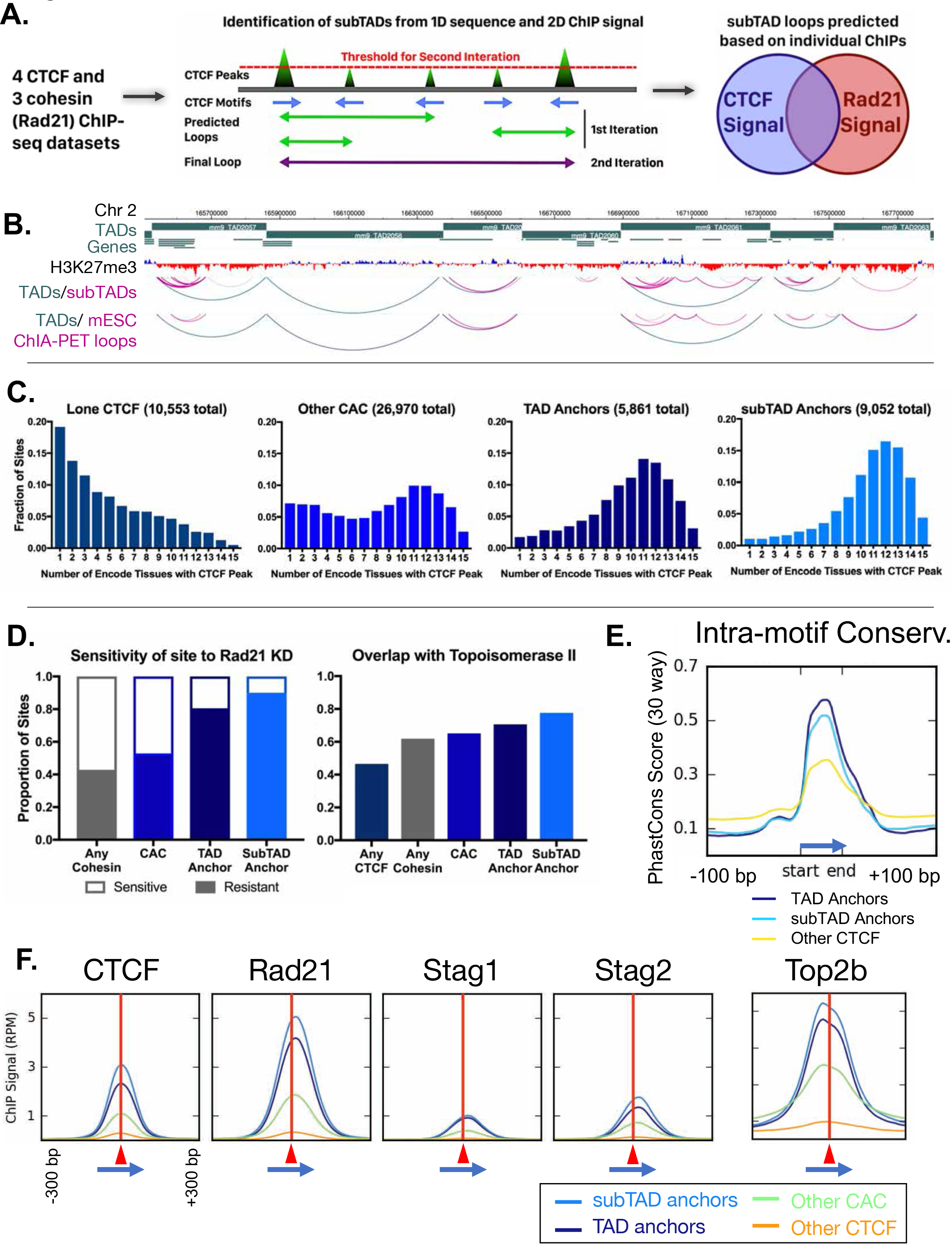
Predicted subTAD loop anchors share many properties of TAD anchors. A. Diagram illustrating subTAD loop prediction using CTCF motif orientation and CTCF and cohesin (Rad21) ChIP-seq binding strength data. Iteration was conducted until 20,000 loops were predicted per sample, prior to filtering and intersection across samples, as detailed in Methods. B. A contiguous 2 Mb section of mouse chromosome 2 showing the relationship between TAD loops (blue) and subTAD loops (pink) in relation to gene structure as well as cohesin interaction loops in mouse ESC identified experimentally by ChIA-PET (5). C. TAD and predicted subTAD loop anchors are more tissue ubiquitous than other categories of CTCF/CAC sites. Each of the four CTCF site subgroups was defined in mouse liver as detailed in Table S1C. The x-axis indicates the number of ENCODE tissues that also have CTCF bound, where a higher value indicates more tissue-ubiquitous CTCF binding. These data are shown for “lone” CTCF binding sites (10,553), non-anchor cohesin-and-CTCF sites (“Other CAC”; 26,970), TAD anchors (5,861), and subTAD anchors (9,052; excluding those at a TAD loop anchor). D. TAD and subTAD anchors are more resistant to the knockdown effects of Rad21+/− haploinsufficency than other CAC sites or cohesin-bound regions. A larger fraction is also bound by the novel extrusion complex factor Top2b. E. Loop anchors show greater intra-motif conservation than other CTCF-bound regions. Shown are the aggregate PhastCons score for oriented core motifs within either TAD (dark blue) or subTAD (light blue) anchors as compared to other CTCF peaks with motifs (yellow). F. Cohesin interacts with the COOH terminus of CTCF, resulting in a shift of ~20 nt in cohesin ChIP signal relative to the CTCF summit (c.f. shift to the right of vertical red line) regardless of category of CTCF binding site (anchor/non-anchor). Blue arrows indicate the CTCF motif orientation and red triangles and vertical lines indicate position of the CTCF signal summit.

#### subTAD loop anchors share many properties of TAD anchors

We examined the set of predicted subTAD loops and their CAC site anchors to investigate their impact on genome structure and gene regulation. For these analyses, we excluded from the subTAD anchor group the 1,632 subTAD anchors that are shared with TAD anchors to ensure that the groups compared are mutually exclusive (Figure S3B). We first sought to determine if the subTAD loop anchors show conserved CTCF binding across multiple ENCODE tissues, as seen for TADs in Figure 1C. Figure 2C shows the tissue distribution of CTCF binding at CTCF binding sites found at subTAD anchors in liver, where an X-axis value of 1 indicates the liver CTCF binding site is occupied by CTCF in only one other tissue, and a value of 15 indicates binding occurs in all 15 mouse tissues where CTCF ChIP-seq data is available. Results show that a large majority of these CTCF binding sites are bound by CTCF in at least 10 of the 15 mouse tissues examined. Indeed, CTCF binding at the TAD-internal subTAD loop anchors is more deeply conserved across mouse tissues than that at TAD boundaries. In contrast, CTCF sites not associated with cohesin binding (lone CTCF sites), and to a lesser extent CAC sites not at subTAD or TAD anchors (other CAC sites) showed much greater tissue specificity for CTCF binding (Figure 2C).

The enrichment of knockdown-resistant cohesin binding sites at TAD boundaries, seen in Figure 1B, may explain the persistence of domain structures following CTCF or cohesin depletion [15, 41]. Further, we found that 80% of TAD anchors and 90% of subTAD anchors are resistant to the loss of cohesin binding in Rad21^+/−^ mice vs. only 52.8% for all CAC sites (Figure 2D). Moreover, a large fraction of TAD and subTAD anchors, 70.6% and 77.6%, respectively, are comprised of “triple sites”, where cohesin and CTCF are co-bound with Top2b, a potential component of the loop extrusion complex [18], vs. only 46.6% for the set of all CTCF sites (Figure 2D). Top2b binding appears to be associated with cohesin binding rather than with CTCF binding, as it is present at enhancer-like CNC sites much more frequently than at CTCF sites in the absence of cohesin (Figure S3C).

TAD and subTAD loop anchors show greater sequence conservation within the core 18 bp CTCF motif than other CTCF sites (Figure 2E). Analysis of sequences surrounding the CTCF core motif did not provide evidence for loop anchor-specific motif usage or cofactor binding (Figure S3D,E). Downstream from the core CTCF motif (within the loop interior) we observed a shoulder of high sequence conservation, as well as additional complex motif usage, likely due to the multivalency of CTCF-DNA interaction, as described in [49]. TAD and subTAD anchors showed broader and more complex CTCF motif usage outside of the core (Figure S3D,E); however, only a small minority of sites contained any specific motif in this flanking region. It is less clear to what extent this broader motif usage is a general property of strongly-bound CTCF regions or of the loop anchors themselves. In fact, we observed a consistent positioning of the cohesin peak just downstream of the CTCF peak, independent of whether the CTCF site was predicted to participate in loop formation or not (i.e. at both loop anchors and “Other CAC” sites; Figure 2F), in accordance with the loop extrusion model and other observations [12, 16, 17].

Given the highly tissue-conserved binding of CTCF at sites that we predicted to serve as subTAD anchors in liver (Figure 2C), we sought direct experimental evidence for the presence of these loops in other tissues. We examined CAC-anchored loops identified in mESC using ChIA-PET [5, 32]. This comparison revealed a striking 68% overlap of our predicted liver subTAD loops with the CAC-anchored loops found in mESC (Figure 2B, Figure S4A). The majority of subTADs are also observed in newer cohesin HiChIP datasets (ChIP followed by Hi-C) [50], and substantial overlap is also observed between high resolution Hi-C loop domains and chromatin contact domains, which tend to be larger in size (Figure S4B,C). Overall ~80% of subTADs are experimentally observed in at least one other tissue. Further, the loops shared between liver and mESC are more robust and showed stronger interactions than mESC-unique loops (Figure S4D).

#### subTAD loops show strong, directional interactions and insulate chromatin marks

TADs are proposed to impact gene expression via two types of insulation: by insulation of chromatin interactions (also called enhancer blocking) and by segregation of chromatin domains, primarily insulation of repressive histone mark spread. We investigated whether subTAD loops demonstrate these dual insulating properties, canonically ascribed to TADs.

CTCF sites were divided into three groups, based on whether they were predicted to anchor TAD loops or subTAD loops, or were not predicted to interact (non-anchor CTCF sites) based on our model. Figure 3A shows the extent to which individual CTCF sites within each group show directional interactions towards the loop interior based on Hi-C data, using a chi-squared metric derived from the same directionality index used to predict TADs genome-wide. This inward bias index quantifies the strength of interaction from a 25 kb bin immediately downstream from the CTCF motif, with a positive sign indicating a downstream (inward) bias with regard to motif orientation, i.e., towards the center of a TAD/subTAD in the case of loop anchors. TAD and subTAD anchors both show strong directional interactions compared to non-anchor CTCF sites, and TAD anchors show stronger interactions than subTAD anchors. For the non-anchor CTCF sites, only those sites containing a strong CTCF motif (FIMO score > 10) were considered, and the predicted directionality was oriented relative to this motif (as there is no left versus right anchor distinction).

**Figure 3:**
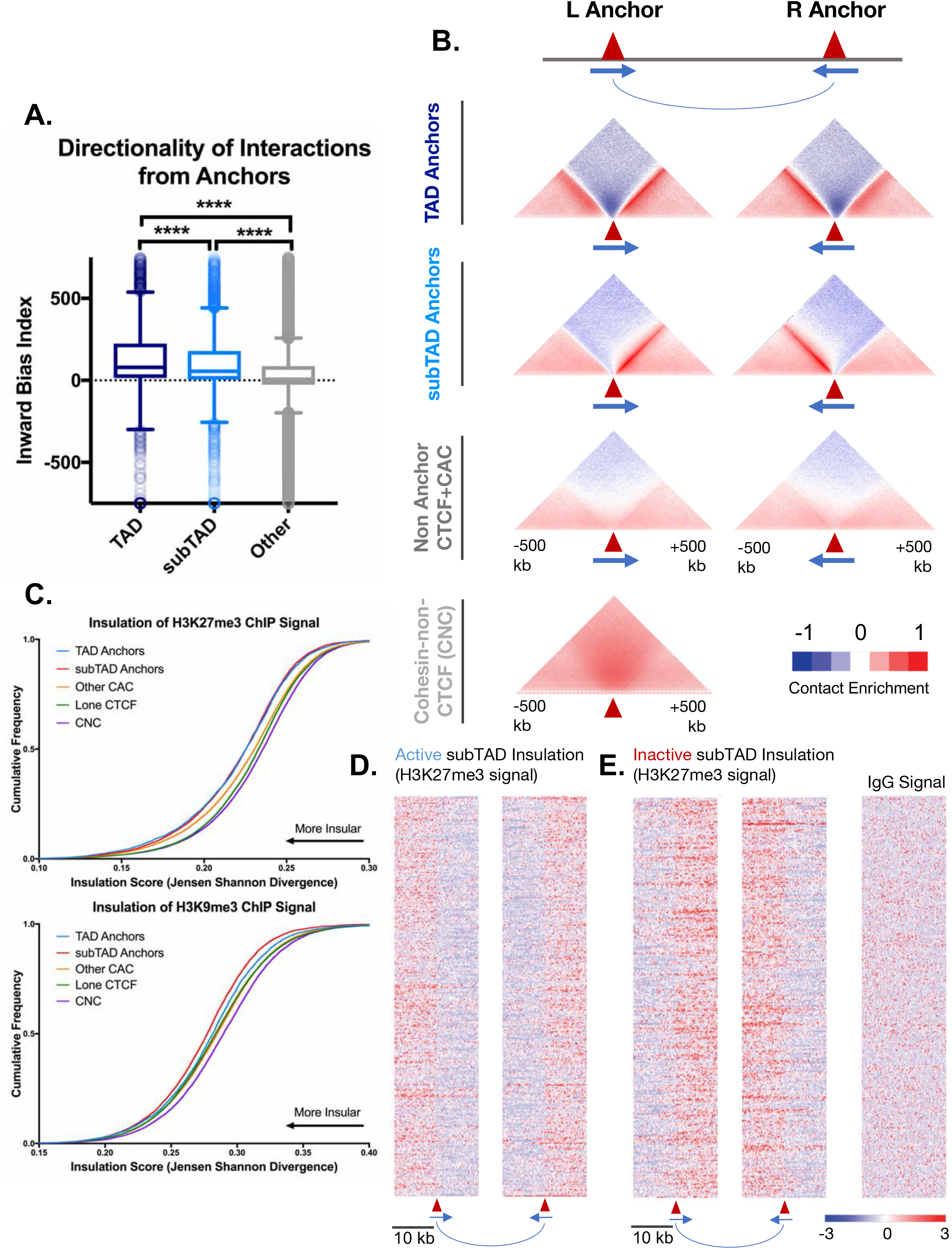
subTADs show directional interactions and insulate chromatin marks. A. TADs and subTADs both show a stronger orientation of interactions downstream of the motif than other CTCF-bound regions. TAD anchors also show higher inward bias than subTADs (p < 0.0001, KS t-test for pairwise comparisons). Inward bias is a chi-square-based metric similar to directionality index but defined on a per peak basis and oriented relative to the motif within the anchor/non-anchor peak. B. In aggregate, TAD and subTAD anchors show stronger and more directionally-biased interactions (contact enrichment, red) than other non-anchor CTCF bound genomic regions. The also show a greater depletion of distal chromatin interactions (contact depletion, blue). TAD anchors also show greater distal contact enrichment with the anchor and more local contact depletion spanning the anchor than subTADs. Red triangles indicate locations of left and right loop anchors and blue arrows indicate CTCF motif orientation. Shading indicates an enrichment (red) or depletion (blue) of contact frequency relative to a genome-wide background model. Figures shown are aggregate plots generated from mouse liver Hi-C data (3) for each set of TAD and subTAD anchors, for the set of non-anchor CTCF sites listed in Table S1C, and for the set of CNC sites as a control (Table S1D). C. TAD and subTAD anchors show greater insulation of the repressive histone marks as measured by Jensen Shannon divergence between H3K27me3 and H3K9me3 ChIP-seq signal upstream and downstream of the anchor region. Shown are JSD values for four classes of mutually exclusive CTCF binding sites (TAD anchors, subTAD anchors, other CAC sites, and CTCF sites lacking cohesin) as well as CNC sites, which are primarily found at enhancers. D. Shown are the top 500 active insulated subTADs, based on high H3K27me3 ChIP-seq signal outside the subTAD (red), and low H3K27me3 signal within the subTAD (blue). Data are expressed as a Z-score of the H3K27me3 signal per bin relative to all H3K27me3 signals within a 20 kb widow centered on all CTCF-bound regions. E. Shown are the top 500 inactive insulated subTADs, based on high signal H3K27me3 signal inside the subTAD (red) and low H3K27me3 signal in neighboring regions (blue). Signal is shown as a Z-score of H3K27me3 signal, as in D.

To visualize features of these anchors, aggregate Hi-C profiles spanning 1 Mb around each anchor were generated for each group of CTCF sites (Figure 3B). Each group was further subdivided into a left (upstream) and a right (downstream) anchor based on its CTCF motif orientation. By aggregating many sites, we can visualize the overall interaction properties of each group of sites at high resolution (5 kb bins), revealing features that are much harder to discern at an individual CTCF site (Figure 3B). TAD and subTAD loop anchors both show strong enrichment of interactions towards the loop interior when compared to CTCF sites that were not predicted to participate in loop formation (non-anchor CTCF sites). Furthermore, subTAD anchors show less enrichment of long-range contacts compared to TAD anchors, likely because of their shorter length compared to TADs. This may also explain the lower inward bias scores of subTAD loops seen in Figure 3A. Depletion of interactions that span across loop anchors (dark blue density above anchor points in Figure 3B) was seen for both TAD and subTAD anchors, however, this local insulation was substantially greater for TAD anchors. This may be due to the compounding impact of adjacent TAD loops, i.e., the end of one TAD is often close to the beginning of an adjacent TAD loop anchor (median distance between TAD anchors = 33.5 kb, Figure S4E). GO term analysis of genes whose TSS fall within these inter-TAD regions revealed enrichment for housekeeping genes (ribosomal, nucleosome, and mitosis-related gene ontologies), whereas neighboring genes found just within the adjacent TADs were enriched for distinct sets of GO terms (Figure S4F; Table S3B, S3C). The nearby but oppositely oriented TAD anchors flanking the inter-TAD regions likely contribute to the more bidirectional interaction pattern for TADs seen in Figure 3B. For instance, the left TAD anchor plot in Figure 3B shows a well-defined pattern of interaction enrichment downstream from the anchor, but also a more diffuse enrichment upstream contributed by upstream loop anchors located at various distances. In contrast, non-anchor CTCF sites do not show strong directional interactions and only very weak distal contact depletion. CNC-bound regions are predominantly found at enhancers [17] and do not show any discernable patterns of insulation or focal and directional interactions (Figure 3B, *bottom*). This demonstrates that the weak contact depletion spanning each of the CTCF-containing groups is likely real, and not an artifact of the background model or other noise.

To determine if subTAD loops share with TADs the ability to block histone mark spread and establish broad, insulated chromatin domains of activity and repression, we analyzed the distribution of two repressive marks in relation to CTCF binding sites: H3K27me3 and H3K9me3. An insulation score based on Jansen Shannon Divergence (JSD) [51] was calculated for ChIP signal distribution within a 20 kb window around each CTCF peak. A low JSD value indicates less divergence from a string representing perfect insulation (i.e., high signal on one side of peak, and low or no signal on the other). This scoring allows for a direct comparison of different classes of CTCF sites, or other TF-bound sites, in terms of their insulation properties. In addition to TAD and subTAD anchors, we examined three other sets of sites as controls: other CAC sites, sites bound by CTCF alone, and CNC sites. Figure 3C shows the cumulative distribution of JSD scores for each set of sites, where a leftward shift indicates greater insulation across the site for the chromatin mark examined. For H3K27me3 signal distribution, TAD and subTAD anchors showed the greatest insulation, but were not significantly different from each other (Kolmogorov-Smirnov (KS) test; p=0.52). The same general trend was observed for H3K9me3 signal insulation; however, subTAD anchors actually showed greater insulation than TAD anchors and all other groups (KS test; p<0.001). CNC sites consistently showed the least insulation of both repressive histone marks, as expected. Sites where CTCF is bound alone showed a small but significant increase in insulation compared to CNC sites, as did the CAC group. As a control, the distribution of IgG signal (input signal) showed much less insulation overall and no significant differences between the various classes of CTCF/cohesin binding sites (Figure S4G).

Figure 3D and 3E show heat map representations`
 of H3K27me3 ChIP-seq signal around the top 500 active and top 500 inactive subTADs, based on a ranked list of JSD insulation scores. These represent subTADs that have significantly lower (or higher) H3K27me3 signal in the loop interior based on the combined rank of JSD insulation scores for each anchor, respectively (p < 0.05, two-sided t-test). For example, Figure 3E shows subTADs with lower H3K27me3 signal within the loop than in neighboring regions (and thus the left anchor transitions from high to low signal, while the right anchor transitions from low to high signal). No such pattern was seen for the IgG (control) ChIP-seq signals for these same regions, indicating this is not an artifact of the sequence read mappability of these regions (Figure S4H).

### III. Impact of subTADs on *cis* regulatory elements in mouse liver

#### Classifying open chromatin regions and defining super-enhancers in mouse liver

TADs, and possibly subTADs, are structurally conserved across tissues, but the activity of enhancers and promoters contained within them is highly tissue-specific [5, 52]. Accordingly, it is important to understand the ability of TADs and subTADs to insulate active enhancer interactions, i.e. enhancer-blocking activity. To address this issue, we examined the distribution of CTCF and cohesin binding sites in relationship to promoters and enhancers across the genome, as well as the impact of TADs and subTADs on their associated genes and enhancers.

We previously identified ~70,000 mouse liver DNase hypersensitive sites (DHS), whose chromatin accessibility is in part determined by TF binding and their flanking histone marks [53, 54]. To assign a function for each DHS, we classified the DHS according to the ratio of two chromatin marks, H3K4me1 and H3K4me3,which are respectively associated with enhancers and promoters [55]. The ~70,000 DHS were grouped into five classes: promoter, weak promoter, enhancer, weak enhancer and insulator (Figure 4A, Figure S5A; see Methods). Promoter-DHS were defined as DHS with a H3K4me3/H3K4me1 ratio ≥ 1.5, and enhancer-DHS by a H3K4me3/H3K4me1 ratio ≤ 0.67 (as first described in [56]). DHS with similar signals for each mark (H3K4me3/H3K4me1 ratio between 0.67 and 1.5) were designated weak promoter-DHS, based on their proximity to TSS and comparatively lower expression of neighboring genes (Figure S5B, S5C). DHS with low signal for both marks were classified as weak enhancers, or as insulators, in those cases where they overlapped a CTCF peak with a comparatively strong ChIP-seq signal (Figure S5A). Using this simplified five DHS class model, we observed that enhancer-DHS bind cohesin largely in the absence of CTCF, while promoter-DHS are bound by both CTCF and cohesin (Figure 4B). Additionally, H3K27ac is enriched at promoter-DHS and enhancer-DHS but not at weak enhancer-DHS, which are less open (lower DNase-seq signal) (Figure 4B) and more distal (Figure S5B). In contrast to enhancer-DHS, insulator-DHS have a well-defined bimodal distribution of tissue-specific vs. tissue-ubiquitous sites based on comparisons across 20 ENCODE tissues (Figure S5D). This supports the proposal that insulators are a unique class of intergenic regions, and not simply enhancers bound by CTCF.

**Figure 4:**
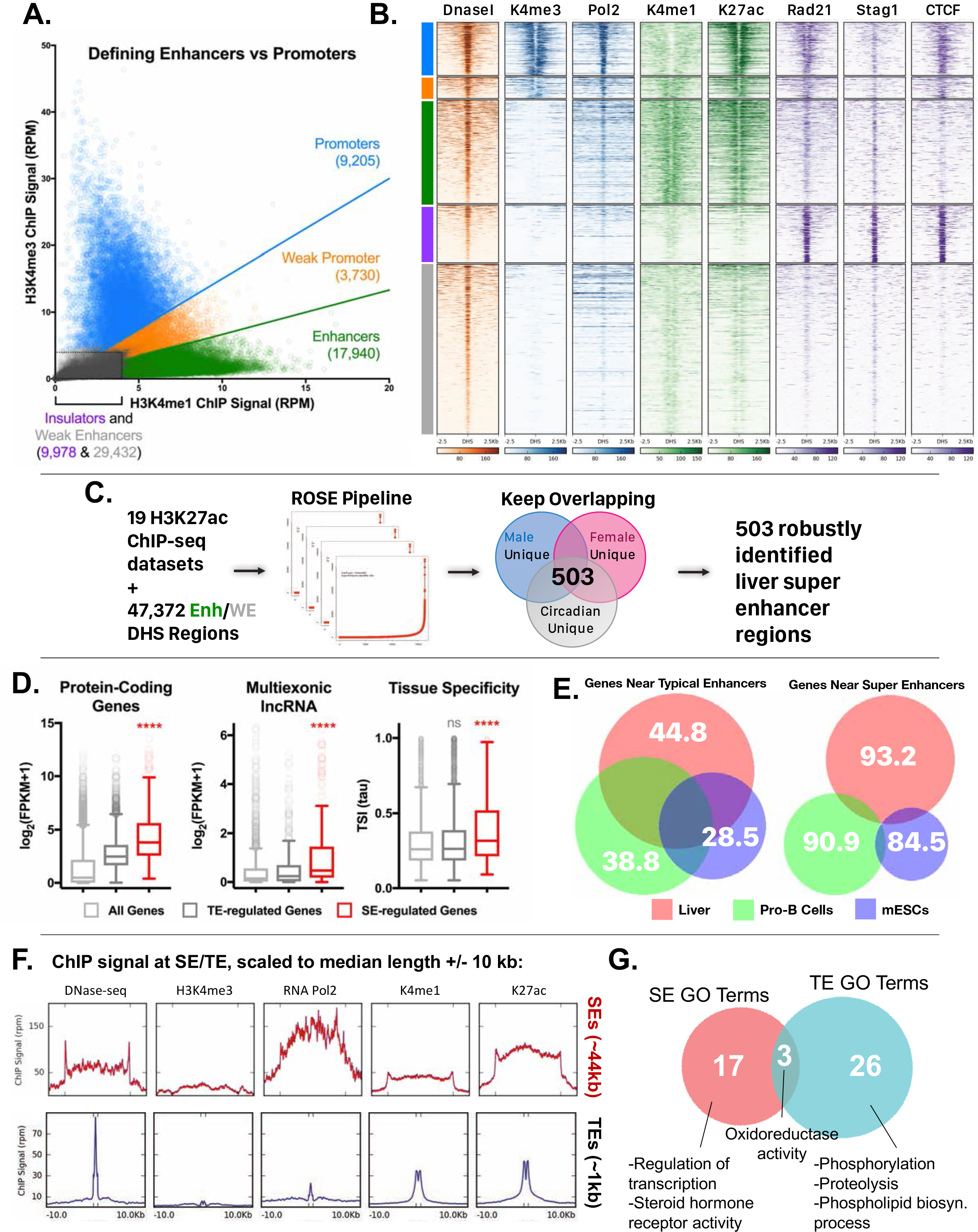
Categorization of cis regulatory elements in mouse liver. A. Classification of open chromatin regions (DHS) in mouse liver, based on relative intensities for a combination of H3K4me1 and H3K4me3 marks, and CTCF ChIP-seq data (see Methods and Figure S5A). Based on the combinatorial signal from these three datasets, five groups of DHS were identified: promoter-DHS, weak promoter-DHS, enhancer-DHS, weak enhancer-DHS, and insulator-DHS. B. A heatmap representation showing that the simplified five-class DHS model shown in panel A captures features such as CNC enrichment at enhancers and K27ac enrichment at enhancers and promoters, with additional features described in Figure S5A. C. Scheme for using 19 published H3K27ac ChIP-seq datasets to identify a core set of 503 superenhancers in mouse liver with the ROSE software package. This core set of 503 super-enhancers was identified in all 19 samples, indicating these super-enhancers are active in both male and female liver, and across multiple circadian time points. D. Genes associated with super-enhancers (SE) are more highly expressed than those associated with typical enhancers (TE), for both protein coding genes and multi-exonic lncRNA genes. Gene expression is presented as log2(FPKM+1) values. Also, these super-enhancer-regulated genes are more tissue specific (higher Tau score) than typical enhancer-regulated genes (KS t-test, p < 0.0001). p-values shown are for pairwise comparisons of SE-regulated genes vs. TE-regulated genes (KS t-test). E. Venn diagrams show the substantial overlap between typical enhancer gene targets across tissues (liver, ESCs, ProB cells), but limited overlap between super-enhancer gene targets (all within 10 kb) for the same tissues. The numbers represent the percent of genes targeted in a given tissue by the indicated class of enhancer (typical enhancers or super-enhancers) that are not targets of the corresponding class of enhancers in the other two tissues. For example, 93.2% of genes targeted by liver super-enhancers are not targeted by the set of super-enhancers identified in either Pro-B or mouse ESCs. Gene targets of each enhancer class were identified by GREAT using default parameters. F. ChIP and DNase-seq signal at typical enhancers and super-enhancers, scaled to their median length (1 kb and 44 kb respectively; indicated by hash marks) with ±10 kb. In addition to a 1.5-2.5-fold higher signal for general enhancer marks, super-enhancers show greater accumulation of RNA polymerase 2, despite no apparent enrichment for the promoter mark H3K4me3. G. Super-enhancers (SE) generally regulate distinct categories of genes than typical enhancers (TE) in mouse liver. While GO terms such as oxidoreductase activity are regulated in common by both classes of enhancers, only super-enhancers are enriched for transcription-regulated terms such as Regulation of Transcription and Steroid hormone receptor activity. Numbers represent the overlap of GO terms (either Molecular Function or Biological Process) in any DAVID annotation cluster (with an enrichment score >1.3) for genes regulated by either typical enhancers or super-enhancers.

A subset of intra-TAD CAC loops are well characterized as insulators of tissue-specific genes with highly active enhancer clusters, termed super-enhancers [5, 57]. To determine if some subTADs correspond to these “super-enhancer domains” we first identified super-enhancers in mouse liver. We used 19 publicly available mouse liver H3K27ac ChIP-seq datasets to score clusters of individual enhancers + weak enhancers DHS identified above (Figure 4C). Super-enhancers were identified separately in male and female liver, as well as in male liver at various circadian time points, to take into account these three key sources of natural variation in mouse liver [53, 58]. In total, we identified 503 core super-enhancers, i.e., super-enhancers that show strong signal regardless of sex or time of day (Figure S6A). Super-enhancers represent 14.1% of all enhancer regions in the genome (6,680 of 47,372 constituent enhancers + weak enhancers), and 2.8% of all enhancer clusters, defined as groups of enhancers within 12.5 kb of one another (503 of all 17,964 enhancer clusters).

Genes neighboring super-enhancers, both protein coding and lncRNA genes, are more highly expressed and tissue-specific when compared to all genes or to all genes neighboring typical enhancers (KS test p-value < 0. 0001; Figure 4D). Consistent with this, only 6.8% of genes proximal to liver super-enhancers are targets of super-enhancers in mESCs or pro-B cells (Figure 4E), whereas 55.2% of genes proximal to liver typical enhancers are proximal to typical enhancers in the other two cell types. Super-enhancers showed much greater accumulation of RNA polymerase 2, despite the lack of the promoter mark H3K4me3 (Figure 4F) and are transcribed to yield eRNAs (Figure S6B). GO terms associated with genes targeted by either typical enhancers or super-enhancers are enriched for liver functions (such as oxidoreductase activity), however, super-enhancer target genes also show enrichment for transcription regulator activity and steroid hormone receptor activity (Figure 4G). These data support the model that super-enhancers drive high expression of select liver-specific genes, including transcriptional regulator genes (Table S2).

Strikingly, 72.2% of core super-enhancers (363/503) are found within a subTAD or within a TAD that contains only a single active gene (FPKM ≥1, and a promoter-DHS within 5 kb of the TSS). By comparison, only 43.6% (17,742/40,692) of typical enhancers are insulated in a similar manner. We also observe an enrichment of single-TSS subTADs (n=3,142) over a random shuffled set (Figure S6C), which could represent tissue-specific genes regulated by super enhancers in other tissues. Genes within these single-TSS subTADs were enriched for ontologies related to transcriptional regulation and phosphorylation (Figure S6D, Table S3D,E). This is consistent with a model of subTADs as functionally inducible units of gene expression, allowing selective transcription in a given tissue or in response to cell signaling events.

To determine the impact of subTADs on the expression of genes with neighboring super-enhancers, we considered two possible gene targets for each super-enhancer, with the requirement that the TSS of each gene target be within 25 kb of one of the individual enhancers that constitute the overall super-enhancer: one gene target is located within the subTAD, and the other gene target crosses a subTAD anchor (Figure 5A, scheme at *top*; genes inside (red) and genes outside (black) of the subTAD). We hypothesized that the true gene target of the super-enhancer will be more highly expressed. Indeed, we found that genes neighboring super-enhancers and found within the same subTAD are significantly more highly expressed than the alternative potential gene targets, located outside of the subTAD (Figure 5A). Similarly, when comparing the tissue specificity of genes within TADs and subTADs to a random shuffled set of subTADs, we observed less variance in the Tau value (index of tissue specificity) for genes within (sub)TAD loops compared to the shuffled set (Figure 5B; KS test p-value < 0.0001). Thus, groups of genes within subTADs are more uniformly tissue specific (as in the case of some super-enhancer-regulated genes) or tissue ubiquitous (as in the case of groups of housekeeping genes).Examples are shown in Figure 5C and 5D, which illustrate the impact of a TAD loop on the expression of *Cebpb* and the impact of a subTAD loop on the expression of *Hnf4a*. The TSS of two other nearby genes, *Ptpn1* and *Ttpa1*, are closer in linear distance to the adjacent super-enhancer than *Cebpb* and *Hnf4a*, respectively, however, any super-enhancer-promoter interactions involving *Ptpn1* and *Ttpa1* would need to cross a TAD or subTAD boundary. In both cases, the genes within the super-enhancer-containing (sub)TAD loop are expressed at least 10-fold higher than genes outside the (sub)TAD loops. Therefore, based on the 3D-structure imposed by these TADs and subTADs, one predicts that the super-enhancers are restricted from interacting with *Ptpn1* or *Ttpa1*, in agreement with their comparatively low expression levels.

**Figure 5:**
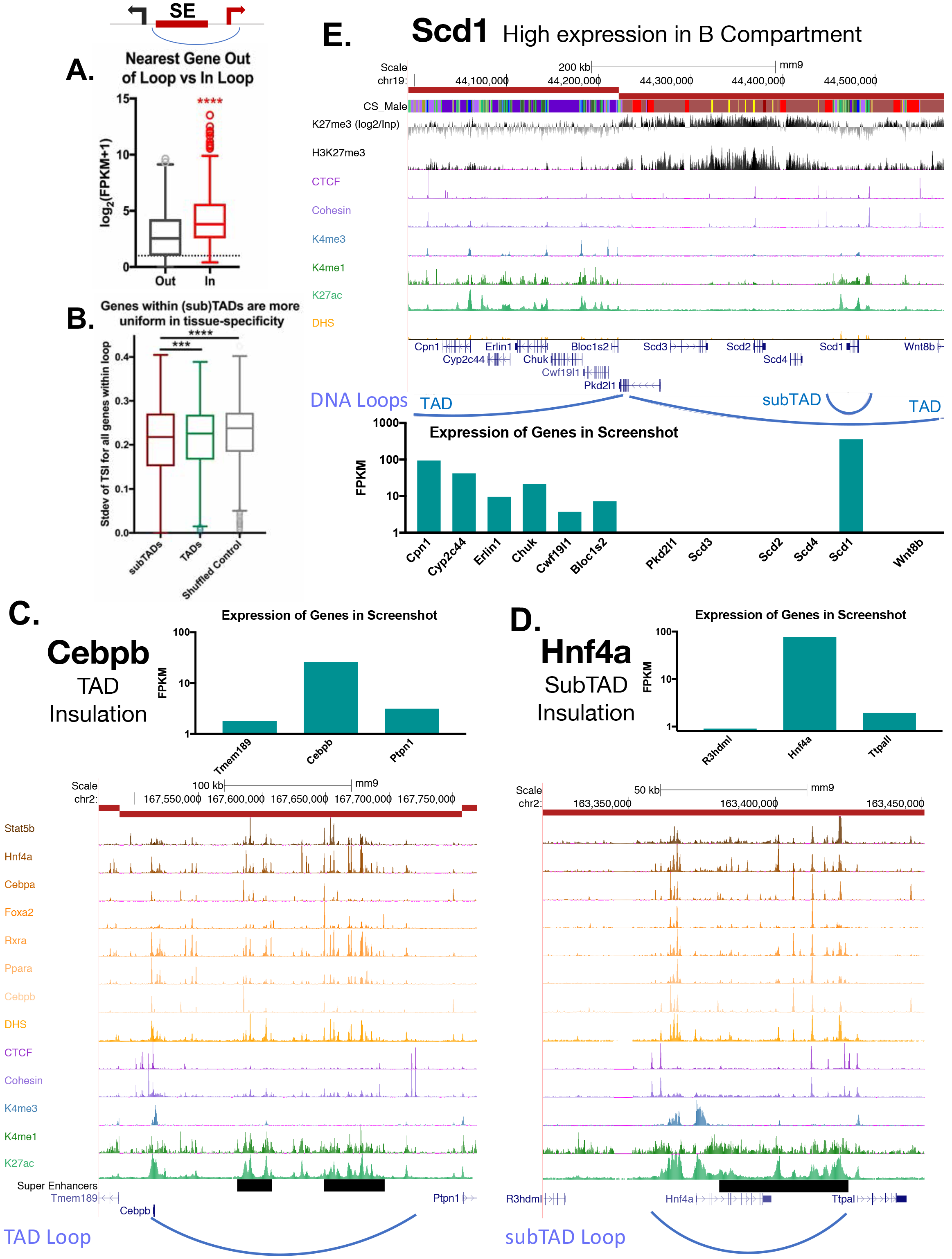
Impact of subTADs on gene expression. A. For those subTADs that include a super-enhancer, two possible gene targets were assigned: one gene within the subTAD and another crossing the subTAD anchor but within 25 kb of the TSS. Box plots show that gene targets within a subTAD are significantly more highly expressed than the alternative, linearly more proximal, gene target. B. Genes within subTADs tend to be more uniformly tissue-specific or tissue-ubiquitous when compared to all genes within TADs or when compared to a shuffled set of random regions matched in size to subTADs. Shown is the standard deviation in Tau values (tissue-specificity index) of genes whose TSS’s are within TADs or subTADs containing at least three TSS. This indicates that sets of three or more genes within subTADs are consistently either more or less tissue specific than random clusters of genes within the same sized genomic spans. C-D. TAD and subTAD loops insulate a subset of super-enhancers (black horizontal bars) with key liver genes, allowing high expression of genes such as the TFs *Cebpb* and *Hnf4a*,relative to their immediate neighbors. *Cebpb* is an example at the TAD scale, while *Hnf4a* shows a subTAD loop. In both cases, the most linearly proximal gene is outside the TAD or subTAD and is expressed at a lower level than the loop-internal genes (and presumptive gene target). E. Insulated subTAD loops allow for expression of select gene targets within otherwise repressed genomic compartments. Shown is a UCSC genome browser screenshot of a transition from an active to a repressed TAD, with the expression of genes within the region shown in a bar graph, below. The obesity-related gene *Scd1* is insulated in a subTAD loop and is the only expressed gene in its TAD (FPKM >100). H3K27me3 marks are shown both as a reads per million signal track (below) and as a signal over an IgG input control (above), expressed as log_2_(H3K27me3 signal / Input signal).

Given the ability subTADs to insulate repressive histone marks (Figure 3C-3E), we considered whether subTAD loops enable high expression of genes within otherwise repressed genomic compartments. As seen in Figure 1G, a minority of genes within inactive TADs are expressed (939 genes expressed at FPKM > 1). The obesity-related gene stearoyl-CoA desaturase-1 (*Scd1*) is one such gene. Figure 5E shows a transitional TAD boundary, with genes in the upstream TAD expressed and associated with low levels of H3K27me3 repressive histone marks. Six genes are in the downstream TAD, but only one of these genes, *Scd1*, is expressed (Figure 5E, *bottom*). The high expression of Scd1 (FPKM > 100) can be explained by its localization in an active subTAD loop that is insulated from the repressive mark H3K27me3 compared to the rest of the TAD. This same structural organization was seen for 291 of the 939 expressed genes found in inactive TADs (Fig. 1G, above), which are contained within subTAD loops. It is unclear what other mechanisms allow for selective expression of the other 648 genes (Table S1E).

#### 4C-seq analysis of super-enhancer contacts at Alb promoter

To test directly for the insulation of a subTAD containing a super-enhancer, we performed 4C-seq analysis for the promoter of albumin (*Alb*), the most highly expressed gene in adult mouse liver. 4C-seq is designed to identify all chromatin contacts originating from a single genomic region (the promoter of *Alb* in this case), known as the 4C viewpoint. Using 4C-seq, we captured many highly specific, reproducible interactions with the *Alb* promoter, a majority of which are localized across an upstream ~50 kb region (Figure 6A, Figure S7A-B). 40% of the interactions are localized within DHS that are constituent enhancers of the *Alb* region super-enhancer. Furthermore, >80% of chromatin contacts are within the *Alb* super-enhancer, and >98% are contained within the *Alb* subTAD loop, in both male and female liver. These interactions become more diffuse with increasing genomic distance from the viewpoint, which may represent dynamic interactions, or alternatively, the effect of averaging across a heterogeneous cell population. Comparing the combined interaction profiles between male and female livers, we observed highly reproducible results, with 92.9% of male interactions also present in female liver (Figure S7C).

**Figure 6:**
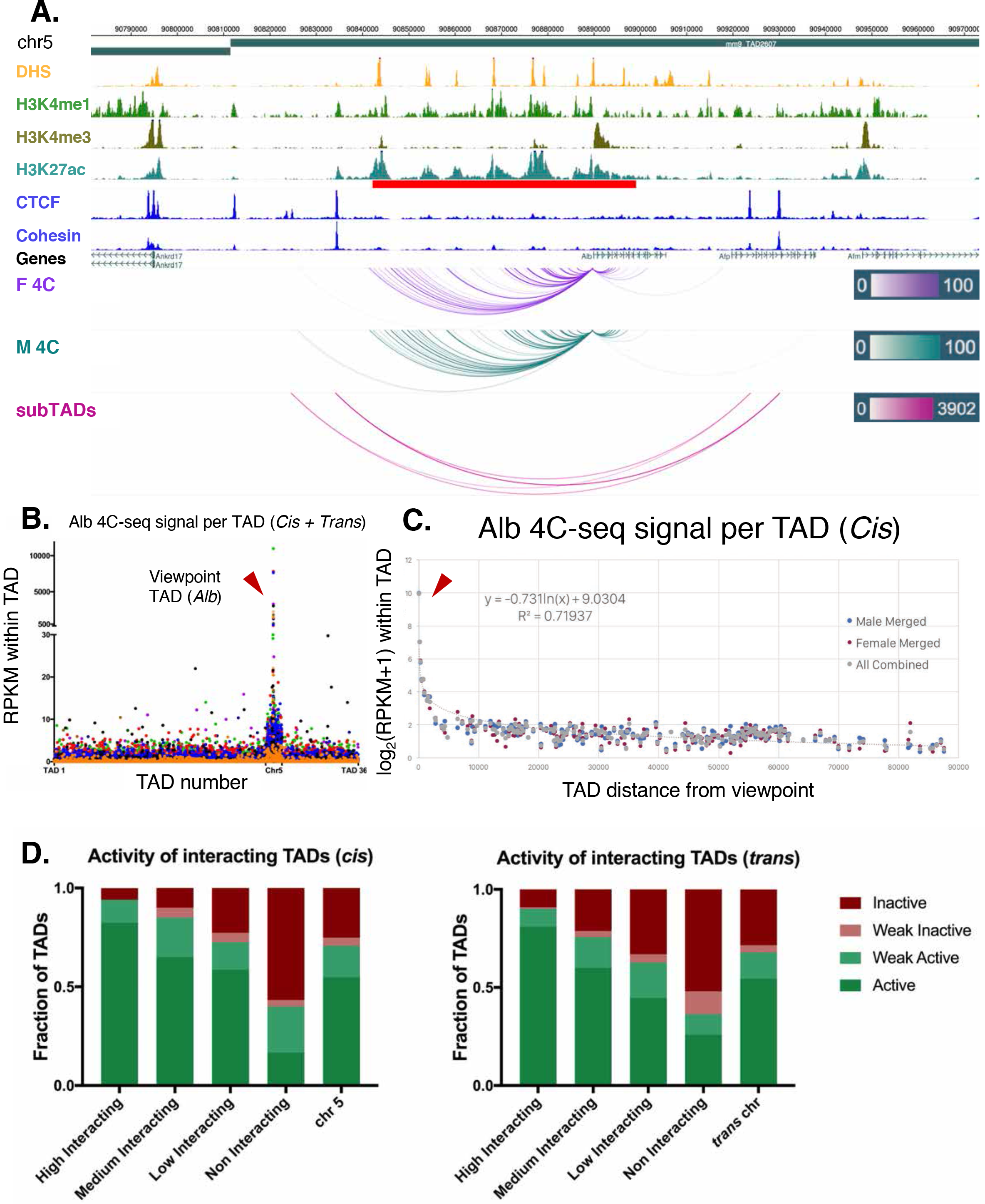
Alb 4C-seq exemplifies subTAD insulation and super-enhancer interaction. A. The *Alb* promoter makes multiple directional contacts with the adjacent super-enhancer region in both male (M) and female (F) mouse liver, as determined by 4C with a viewpoint at the *Alb* promoter. All reproducible interactions occur within the TAD loop containing the *Alb* TSS and its super-enhancer (red), and all but two contacts in male liver occur within the predicted subTAD loop (pink). 4C-seq interaction scores are shown as −log10(pval) across replicates as calculated by R3C-seq (see Methods). Also see Figure S7. B. The 4C interaction signal within the *Alb* TAD is orders of magnitude above the background signal and generally decays with distance. Far-*cis* and *trans* interactions are represented on a per TAD basis, expressed as RPKM per TAD, to control for sequencing depth and TAD length. The overall background within chromosome 5 is significantly higher than all *trans* chromosomes; immediately adjacent TADs also show higher 4C signal than the overall cis background. The 4C signal decayed to background levels within approximately 3 TADs of the *Alb* viewpoint TAD. Each data point represents a single TAD and each color represents a 4C-seq replicate. C. Background model used for distal *cis* interactions, showing a rapid decay in per TAD signal intensity. Each data point represents a single TAD along chromosome 5. D. Distal *cis* and *trans* TADs that highly interact with the *Alb* promoter tend to be active TADs, while a majority of TADs that interact less than the background model are predicted to be inactive. Along the *cis* chromosome, a simple inverse logarithmic decay of signal per TAD was used to determine the background, while the 4Cker package was used to determine high, medium, low, and non- interacting TADs in *trans* based on a hidden markov model with adaptive windows better suited for low signal regions.

Looking beyond local interactions, we observed interaction frequencies that span several orders of magnitude going from local (intraTAD) to *cis* (intra-chromosomal) and *trans* (genome-wide) interactions. Thus, for the *Alb* promoter, the TAD 4C signal per TAD was up to >1000 RPKM for local interactions, >100 RPKM for *cis* interactions within 3 TADs of the viewpoint, and ~10 RPKM beyond that. *Trans* interactions were almost exclusively < 10 RPKM (Figure 6B, 6C). Using a separate background model for far-cis and *trans* 4C signals, we categorized TADs as either high, medium, low, or non-interacting (see Methods). As *Alb* is the most highly expressed gene in liver and is proximal to a strong super-enhancer, we expected that distal interacting regions would also be active genomic regions, as proposed in the transcription factory model of nuclear compartmentalization [59]. Indeed, we found that >80% of the distal high interacting TADs (both far-cis and *trans*) were active, and >90% were either active or weakly active. In contrast, only 16.6% of the non-interacting TADs in *cis* and 25.9% of those in *trans* were considered active (Figure 6D). Furthermore, genes in the interacting regions are more highly expressed than genes in *Alb* 4C non-interacting regions, and the vast majority are found in active TADs (Figure 1E), as determined by analysis of the Hi-C data alone (Figure S7D,E).

## DISCUSSION

We present a computational method that uses 2D (ChIP-seq) and 1D (DNA sequence) information to predict 3D-looped intra-TAD structures anchored by cohesin and CTCF (CAC sites), and we provide evidence that the subTAD loops predicted underpin a general mechanism to constrain the interactions of distal enhancers to specific gene targets. While select instances of CAC-mediated loop insulation within TADs have been described [5, 60, 61], our work establishes that this phenomenon is a more general feature of genomic organization and regulation than previously appreciated. The subTADs described here are nested, CAC-mediated loops whose formation may be a result of extrusion complex pausing within larger domains (i.e., TADs). Their ultimate impact is likely to constrain the available promoter contacts for a given distal enhancer, or vice versa [35]. While the precise mechanism that differentiates TAD- and subTAD-forming CTCF sites (anchors) from other CTCF binding sites (non-anchors) in the genome is unknown, we provide evidence that loop-forming CTCF sites at TAD and subTAD anchors, but not other CTCF site, are highly insular. This difference in insulation is apparent from the blockage of repressive histone mark spread and by the inhibition of chromatin contacts across subTAD and TAD boundaries. The impact of this insulation is highlighted for super-enhancer regions, such as the superenhancer upstream of *Alb*,where local insulation by CAC-anchored subTAD loops both enables and constrains strong near-*cis* interactions, which enables high expression of *Alb* and presumably also other liver-expressed genes regulated by super-enhancers. Weaker *trans* interactions with distal active regions were also observed, and are likely driven by a distinct mechanism, such as aggregation of transcription factories or super-enhancers [26, 62].

Genomic interactions occur at three levels: (1) compartmentalization, where inactive regions localize to the nuclear periphery and active chromatin compartments aggregate toward the center of the nucleus in *cis* or *trans* in a largely cohesin-independent manner, as proposed in the transcription factory model [7, 12, 41, 62]; (2) CAC-dependent looping, which generates mostly tissue-invariant scaffolds along the linear chromosome [1, 13, 35]; and (3) enhancer-promoter looping within CAC-loops, which may be directed by cohesin non-CTCF (CNC) sites, mediator, or tissue-specific TFs [5, 16, 17]. If TADs define the broad domain in which a cohesin-driven extrusion complex generally operates, we have shown a simple method to identify loops within this region that form as a result. We have used the term subTAD to describe these loops to highlight their intra-TAD range, although they are functionally similar to loop domains, isolated cliques, and insulated neighborhoods, which tend to overlap or be contained within TADs [12, 13, 35]. It should be noted that our approach cannot predict CTCF-independent loops such as those mediated cohesin alone (enhancer-promoter loops), although such loops are likely constrained by CAC driven subTADs, as was highlighted by our Albumin 4C-seq results.

The method for subTAD identification described here builds on the preference for inward-facing CTCF motifs evident from high resolution Hi-C data [12, 13], and will be highly useful for the identification of intra-TAD CAC loops for the large number of cell lines and tissues that lack high resolution Hi-C data. In these cases, intra-TAD loop domains cannot be identified because there is not sufficient local enrichment to calculate a corner score with the arrowhead algorithm [12]. Further, while we used liver Hi-C data to filter out longer CAC loops, the general conservation of TADs across both tissues and species [1, 3] broadens the applicability of our method to other organisms with well-annotated genomes, even when Hi-C data is not available and TAD boundaries have not been determined. Thus, even in the absence of TAD coordinates, our method identifies TAD and subTAD looping events, which provides an invaluable first approximation for understudied organisms. As we have tuned our parameters to identify loop structures comparable in size and number to those found previously in mouse models, the exact cutoff values may need to be adjusted for other model organisms.

The identification of subTADs at high resolution from Hi-C data requires extremely deep sequencing of Hi-C libraries, something that has only been achieved in a small number of cell lines [12], and is costly, both in terms of computational and experimental laboratory resources. Alternative strategies have been proposed to reduce the need for extreme deep sequencing to identify interactions at high resolution [36, 63]. While these approaches are beginning to make 5-10 kb resolution possible, the sequencing depth and cost will likely remain out of reach for many labs. The computational method described here considers both CTCF and cohesin peak strength as the primary predictor of subTAD loop strength, which is a reasonably good predictor of interactions [13, 31]. Antibody enrichment for select genomic regions followed by chromosome conformation capture, as implemented in ChIA-PET, is a common experimental alternative, and can identify “many to many” interactions, instead of “all to all” interactions identified by Hi-C [33]. Of note, ChIA-PET and other 3C-based antibody enrichment methods select for genomic regions that are highly bound by the protein(s) of interest (e.g., CTCF and cohesin), making it difficult to differentiate strength of anchor protein binding from strength of chromatin interaction between the anchors. Of the various CTCF loops described in the literature, subTADs are most similar to insulated neighborhoods, which are proposed to rectify the observation of smaller and more abundant loops from ChIA-PET experiments with the established TAD model of large loops from Hi-C experiments [5, 12, 38]. Of practical importance, ChIA-PET requires ~10-fold more extensive deep sequencing per sample (~400 million reads) than is needed to obtain the CTCF and cohesin ChIP-seq data utilized in our computational analysis to identify subTAD loops.

We found that TAD and subTAD loop anchors together comprise 27% of all liver CTCF binding sites, consistent with the 30% of murine ESC CTCF peaks that overlap insulated neighborhood anchor regions [35]. It is unclear what role the remaining, typically weaker, 70% of CTCF sites play in organizing the nucleus. Some of these non-(sub)TAD anchor CTCF sites may serve other, unrelated functions, given the ability of CTCF to interact with other TFs, bind RNA, and regulate splicing mechanics [64–66]. Alternatively, some of these CTCF sites may anchor loops present in only a minority of cells in the population analyzed, which would account for their overall weaker signals. Indeed, early single cell Hi-C experiments suggested that TADs are present in individual cells [67], however, more recent studies indicate greater cell-to-cell variability in TADs than was previously appreciated, although the presence of distinct active and inactive genomic compartments is consistently seen across most cells [40, 68]. Truly high-resolution elucidation of single cell intra-TAD structures may not be possible due to the intrinsic limitation of two potential ligation events per fragment in any given cell.

Regarding the genomic distributions of cohesin and CTCF binding sites, we found that CNC sites are primarily present at enhancers and weak enhancers, while CAC sites are found at insulators and also at promoters, which we defined as DNase hypersensitive sites (DHS) with high a histone-H3 K4me3/K4me1 ratio. Others find that promoters, when defined as the set of all TSS upstream regions, including those not at a DHS, are bound by cohesin alone [16, 17]. Further, we found that CTCF-bound open chromatin regions distal to promoters (insulator-DHS) show features that distinguish them from other classes of open chromatin (promoter-DHS and enhancer-DHS), including the absence of enhancer marks and their general conservation across tissues. Thus, these insulator-DHS regions are not simply enhancers with CTCF bound. Supporting this, insulator regions consistently show less intrinsic enhancer activity than weak enhancers in *in vivo* enhancer screens [69]. It is less clear what role CTCF binding in the absence of cohesin plays in the nucleus, as we found such sites lack insulating activity and also lack strong directional interactions. Since the binding of CTCF is always intrinsically directional, due to its non-palindromic motif, the absence of directional interactions from CTCF-non-cohesin sites suggests that the directionality of interactions with CTCF sites at TAD and subTAD loop anchors is conferred by additional factors, such as cohesin as a part of the extrusion complex [11, 13].

Top2b is another factor that is proposed to be a key contributor to loop extrusion complexes [18]. However, our findings indicate that the interactions of Top2b involve binding to cohesin, and not CTCF, as suggested by the high frequency of CNC sites bound by Top2b vs. very low frequency of binding to CTCF-non-cohesin sites (Figure S3C). Furthermore, binding by Top2b does not distinguish TAD from subTAD loop anchors. Indeed, by all metrics tested, we found no TF or motif that differentiates TAD from subTAD loop anchors, although the existence of some unknown differentiating factor cannot be ruled out. Cohesin has the ability to stabilize large protein complexes, as indicated by its role in stabilizing clusters comprised of up to 10 distinct TFs at enhancers [17], and could thus facilitate the binding of additional unknown proteins to the loop extrusion complex.

Cohesin is continuously recycled throughout the genome by loading and release factors [47], and so it is unclear how insulator activity is effectively maintained at TAD and subTAD anchors in such a dynamic environment. We found that CNC sites, which are primarily at enhancers, consistently show the least insulation of repressive histone marks, just as they show the least insulation of chromatin contacts. This provides further evidence that TAD and subTAD loop anchors are functionally unique sites, and are not a moonlighting feature of CTCF bound to enhancer regions. Furthermore, while enhancers are strongly enriched for genetic non-coding variants, genetic variations at loop anchors are rare [35]. Mutations at loop anchors can result in dramatic phenotypes like polydactyly or tumorigenesis [70] and often occur in cancer [28]. Selective disruption of CAC-mediated loop anchors using genomic editing tools results in aberrant chromatin contacts and misregulation of neighboring genes in a largely predictable manner, although the extent of redundancy is not always clear when multiple anchors are present [5, 35, 60].

The computational method for subTAD loop discovery, presented here, is a substantial improvement over prior implementations of computational loop prediction [13, 31]. Thus, the loops we identified were longer and fewer in number (~9,500 vs 60,000), showed much stronger insulation of chromatin interactions and greater insulation of repressive histone marks, and displayed considerably greater overlap with cohesin-mediated loops identified by ChIA-PET (Figure S2). Key features of our computational method include the consideration of both CTCF and cohesin binding strength, TAD structure and consistency across biological replicates. Our use of both CTCF and cohesin binding strength in predicting subTAD loops is supported by a recent study of CTCF sites nearby the mouse α-globin gene cluster, where the presence of CTCF alone was not sufficient to predict DNA loop interactions, and where insulation by individual CAC sites ranged widely - from none to moderate to very strong insulation - in direct proportion to the strength of CTCF binding, as revealed by deletion of individual CTCF sites [61]. Furthermore, we developed a simple extension of our computational method to predict TAD anchors, and thereby overcome the limitation of the low resolution of standard depth Hi-C datasets in identifying TAD boundaries. We could thus identify well-defined inter-TAD regions, which were enriched for unique gene ontologies, notably, housekeeping genes with ribosomal, nucleosome, and mitosis-related functions. A further extension of our findings would be the explicit use of the TAD, subTAD and inter-TAD loop coordinates defined here to improve gene target assignments for distal regulatory elements, based on the insulating capacity of these looped domains. Such an approach will be beneficial for the many model systems where distal enhancer activity is the clear driver of tissue specificity or a given disease state [35]. The ability to identify subTAD loops based solely on CTCF motif orientation and CTCF and cohesin ChIP-seq binding data, and then use these loops to improve gene target assignments for distal regulatory elements is likely to constitute a substantial improvement over “nearest gene” or other more nuanced target assignment algorithms, such as GREAT [71].

In conclusion, our studies reveal that while TAD structures are readily apparent in routine Hi-C experiments, and thus are better characterized than internal TAD sub-loops, their structural organization and functional impact on the genome is not unique. Thus, the 9,543 subTAD loops that we identified have strong intra-TAD cohesin-and-CTCF-bound anchor regions. Structurally, these subTADs appear to be formed by the same loop extrusion mechanism that is responsible for TAD formation. Functionally, we hypothesize that these subTADs contribute to nuclear architecture as intra-TAD scaffolds that further constrain enhancer-promoter interactions. Further, our findings reveal that subTADs maintain key properties of TADs, most notably insulation of interactions and insulation of repressive histone marks. The insulation provided by subTADs may enable high expression of super-enhancer target genes, as illustrated for *Alb*,as well as high expression of individual genes within otherwise inactive TADs, as exemplified by *Scd1* and the many other single gene subTADs that we identified. Given the increasing interest in interactions of genes with distal enhancers and other intergenic sequences, the rapid and cost-effective method described here for identification of subTAD structures that constrain these long-range interactions may prove invaluable in many areas of research.

## MATERIAL AND METHODS

### Animals and sample processing

Adult male and female CD-1 mice (ICR strain) were purchased from Charles River Laboratories (Wilmington, MA) and were housed in the Boston University Laboratory Animal Care Facility. Animals were treated using protocols specifically reviewed for ethics and approved by Boston University’s Institutional Animal Care and Use Committee (IACUC). Livers were collected from 8-week-old mice euthanized by cervical dislocation. Livers were harvested, rinsed in cold PBS, and homogenized with a Potter-Elvehjem homogenizer using high sucrose homogenization buffer (10 mM HEPES (pH 7.5), 25 mM KCl, 1 mM EDTA, 2 M sucrose, 10% glycerol, 0.05 mM DTT, 1 mM PMSF, 0.15 mM spermine, 0.2% (v/v) spermidine, 1 mM Na orthovanadate, 10 mM NaF, and Roche Complete Protease Inhibitor Cocktail) to prevent aggregation of nuclei and preserve chromatin structure. The resulting slurry was transferred on top of a 3 ml cushion of homogenization buffer followed by ultracentrifugation at 25,000 RPM for 30 min at 4°C in an SW41 Ti rotor to remove cellular debris. The supernatant was carefully decanted to remove liquid, and residual solid debris on the tube walls was removed with a sterile spatula and a dampened Kimwipe. Nuclei were then resuspended in 1 ml of crosslinking buffer (10 mM HEPES buffer (pH 7.6), 25 mM KCl, 0.15 mM 2-mercaptoethanol, 0.34 M sucrose, 2 mM MgCh) and transferred to a 1.5 ml Eppendorf tube. To ensure consistent crosslinking, tubes were incubated for 3 min at 30°C prior to the addition of formaldehyde to a final concentration of 0.8% (v/v). Samples were incubated in a 30°C water bath for 9 min with periodic mixing. Crosslinking was halted by the addition of 110 ul of 1 M glycine, followed by a 5 min incubation at room temperature. The crosslinked material was layered on top of 3 ml of high sucrose homogenization buffer and then centrifuged as above. The crosslinked nuclear pellet was resuspended at 4°C in 1 ml of 1× Radioimmunoprecipitation assay (RIPA) buffer (50 mM Tris-HCl, 150 mM NaCl, 1% IPEGAL, 0.5% deoxycholic acid) containing 0.5% SDS and protease inhibitors until homogenous.

### Sonication

Crosslinked nuclei in RIPA buffer containing 0.5% SDS were transferred to 15 ml polystyrene tubes (BD Falcon # 352095) for sonication using a Bioruptor Twin instrument (UCD-400) according to the manufacturer’s instructions. Briefly, samples were sonicated at 4°C for 30 sec ON and 30 sec OFF at high intensity for a total of 75 cycles. Sonicated material was transferred to 1.5 ml Eppendorf tubes, and large debris was cleared by centrifugation at 18,000 × g for 10 min at 4°C. The bulk of this material was snap frozen in liquid nitrogen and stored at −80°C for immunoprecipitation, except that a small aliquot (15 ul) was removed to quantify material and ensure quality by gel electrophoresis, as follows. Aliquots from each sample were adjusted to 0.2 M NaCl, final concentration, then incubated for 6 h at 65°C. After a three-fold dilution in nuclease-free water, 5 ug of RNase A (Novagen: #70856) was added and samples were incubated for 30 min at 37°C. Samples were then incubated for 2 h at 56°C with 20 ug of Proteinase K (Bioline; BIO-37084). This material was then quantified in a dilution series using PicoGreen assay (Quanti-iT dsDNA Assay Kit, broad range, Invitrogen) and analyzed on a 1% agarose gel to ensure the bulk of material was within 100-400 bp.

### Immunoprecipitation

Immunoprecipitation and downstream steps were as described previously [53]. Protein A Dynabeads (30◻ul; Invitrogen: 1002D) were incubated in blocking solution (0.5% BSA/PBS) with 5 ul of antibody to CTCF (Millipore #07-779) or Rad21 (Abcam #992) for 3 h at 4°C. As a control, 1 ul rabbit IgG was used (Santa Cruz: sc-2027). Bead immune-complexes were washed with blocking solution, followed by overnight incubation with 70 ug of chromatin. After washing with 1× RIPA (containing 0.1% SDS) and reverse crosslinking as above, DNA was purified using the QIAquick Gel Extraction Kit (Qiagen #28706). Purified DNA was quantified using Qubit (Invitrogen DNA HS# Q32854) with pulldown ranging from 1-25 ng.

### Sequencing

ChIP libraries were prepared for sequencing using NEBNext Ultra II DNA Library Prep Kit for Illumina according to the manufacturer’s instructions (NEB, cat. #E7645). All samples were subjected to doublesided SPRI size selection prior to PCR amplification (Agencourt AMPure XP; Beckman Coulter: A63882). Samples were assigned unique barcodes for multiplexing, and amplified for 8 rounds of PCR amplification with barcoded primers (NEB, cat. #E7335). Samples were sequenced either on an Illumina Hi-Seq 4000 instrument at the Duke Sequencing Core or an Illumina Hi-seq 2000 instrument at the MIT BioMicroCenter, giving 50 bp single end reads at a depth of ~11-19 million reads per sample. A total of four CTCF and three Rad21 (cohesin) ChIP-seq samples were analyzed, representing four male mouse livers. CTCF sequencing sample G133_M9 did not have a matching cohesin ChIP-seq dataset from the same liver sample, and thus was matched to a merged sample from all three cohesin ChIP replicates (merged at the fastq file level, with processing described below). Raw and processed sequencing data are available at GEO accession number: GSE102997.

### General ChIP-seq analysis pipeline

Sequence reads were demultiplexed and mapped to the mouse genome (build mm9) using Bowtie2 (version 2.2.9), allowing only uniquely mapped reads. Peaks of sequencing reads were called using MACS2 (version 2.1.1) to identify regions of high signal over background. Peaks were filtered to remove blacklisted genomic regions (www.sites.google.com/site/anshulkundaje/projects/blacklists) and also regions called as peaks that contain only PCR duplicated reads, defined as >5 identical sequence reads that do not overlap any other reads. All BigWig tracks in screenshots are normalized for sequencing depth expressed as reads per million mapped reads (RPM) using Deeptools (version 2.3.3). Unless otherwise indicated all pairwise comparisons in Figures were performed using a Kolmogorov-Smirnov test, where **** indicates p ≤ 0.0001, *** p ≤ 0.001, ** p ≤ 0.01, or * p ≤ 0.05.

### Motif Finding

Motif analysis within CTCF peak regions was performed using the MEME Suite (version 4.10.0; FIMO and MEME-ChIP options). FIMO was used to assign CTCF motif orientation and motif scores for CAC sites and to discover individual motif occurrences. *De novo* motif discovery was carried out using MEME-ChIP using default parameters (Figure S3D,E). Similar results were obtained using Homer (version 4.8). Alternative CTCF motifs were downloaded from the CTCFBSDB 2.0 (http://insulatordb.uthsc.edu/download/CTCFBSDB_PWM.mat), however, these did not substantially change any results performed using the core JASPAR motif (MA0139.1). These motifs were explicitly used in Figure S3E, where no difference between subTAD and TAD anchor motif usage was observed.

### 4C-seq protocol

Four male and four female mouse livers were processed for *Albumin*-anchored 4C-seq analysis using published protocols, with some changes for primary tissue [72]. To adapt the protocol for liver, care was taken to rapidly isolate single cell or nuclei suspensions prior to crosslinking. Specifically, two approaches to crosslinking were taken and both gave similar results. One male and one female mouse liver sample was processed through the crosslinking step as described for the ChIP protocol, above, prior to quantification of nuclei. The other samples (3 males and 3 females) were crosslinked as follows. Half of a liver (~0.5 g) was dissected from each mouse, the gall bladder was removed, and the liver was rinsed with PBS. The liver was then minced and rapidly processed with 10 strokes with a Dounce homogenizer in PBS containing protease inhibitors (PBS-PI; 1X Roche Complete Protease Inhibitor Cocktail; Roche #11697498001). The resulting slurry was passed through a 40-micron cell strainer (Corning #431750), then pelleted and rinsed with PBS-PI (centrifugation at 1,300 RPM for 5 min at 4°C). Following an additional spin, the cell pellet was resuspended well in 9 ml of PBS-PI at room temperature. 270 ul of 37% formaldehyde was added to give a final concentration of 1%, and crosslinking was carried out for 10 min with nutation at room temperature. The remaining formaldehyde was quenched with 1.25 ml of 1 M glycine. Crosslinked cells were pelleted and rinsed with PBS twice (as above) prior to lysis. The supernatant was removed following the second wash, and cell pellets were resuspended in 8 ml of lysis buffer and incubated on ice for 40 min with occasional mixing (50 mM Tris (pH 7.5), 150 mM NaCl, 5 mM EDTA, 0.5% NP-40, 1% TX-100, and 1X Complete Protease Inhibitor Cocktail). Lysed cells were spun down at 2,000 RPM for 5 min at 4°C then washed twice with PBS-PI, as above. Nuclei were pelleted, quantified using an Invitrogen Countess instrument, and snap frozen in 10 million nuclei aliquots. Primary digestion of 10 million nuclei with 50,000 U of DpnII (NEB: #R0543) was performed overnight at 37°C in 450 ul of NEBuffer 3 (NEB: #B7003S; 100 mM NaCl, 50 mM Tris-HCl, 10 mM MgCl_2_, 1 mM DTT, pH 7.9) with agitation at 900 RPM. After confirming digestion by agarose gel electrophoresis, DpnII was inactivated with SDS (2%, final concentration) and the samples then diluted 5-fold in 1× ligation buffer (Enzymatics #B6030). 200 U of T4 DNA ligase was added and primary ligation was carried out overnight at 16°C (Enzymatics #L6030). Ligation was confirmed by analysis of a small aliquot on an agarose gel, and reverse crosslinking was conducted by overnight incubation with 600 ug proteinase K at 65°C. After RNase A digestion and phenol/chloroform cleanup, samples underwent secondary digestion with 50 U of Csp6I (Fermentas #ER0211) overnight at 37°C in 500 ul of 1× Buffer B (Fermentas: #BB5; 10 mM Tris-HCl (pH 7.5), 10 mM MgCl_2_, 0.1 mg/ml BSA). Csp6I was then heat inactivated for 30 min at 65°C. Samples were diluted 10-fold and secondary ligation was carried out as above, overnight at 16°C. The final PCR template was purified by phenol/chloroform clean up, followed by QiaPrep 2.0 column cleanup (Qiagen #27115). PCR reactions were performed using inversely-oriented 4C primers specific to the *Alb* promoter (sequences shown in bold, below) with dangling 5′ half adapter sequences (Reading primer:ACACTCTTTCCCTACACGACGCTCTTCCGATCT**GGTAAGTATGGTTAATGATC**; Non-reading primer: GACTGGAGTTCAGACGTGTGCTCTTCCGATCT**CTCTTTGTCTCCCATTTGAG**). This design has two advantages: 1) the addition of barcodes in a secondary reaction allows a primer to be reused across samples; and 2) it avoids barcoding at the start of 4C read, which would reduce the mappable read length available for downstream analyses. 4C templates were amplified using Platinum Taq DNA polymerase (Invitrogen #10966026) under the following conditions: 94°C for 2 min, 25 cycles at (94°C 30 s, 55°C 30 s, 72°C 3 min), then 4°C hold. A total of eight samples were analyzed (four males, M1-M4; and four females, F1-F4). For liver samples M3, M4, F3 and F4, eight identical PCR reactions processed in parallel were pooled to limit the impact of PCR domination in any single reaction. For liver samples M1, M2, F1 and F2, two PCR reactions for each sample were sequenced separately then pooled at the fastq file level for downstream analyses. We observed that the 8 PCR pool reactions gave a more reproducible profile than single PCR reactions. After pooling, 4C samples were purified using AMPure XP beads (Beckman Coulter: #A63882) at a 1.5:1 ratio of beads to sample, washed with 75% ethanol, dried, and resuspended in 0.1× TE buffer to elute the DNA. 4C-seq samples were multiplexed and PCR amplified using standard NEB barcoded primers (NEB #E7335) as was done for ChIP-seq library preparations, but for five additional PCR cycles (total of 30 cycles of PCR per sample; 25 cycles with viewpoint primers followed by 5 cycles with barcoded NEB primers). 4C libraries were sequenced on an Illumina 2500 instrument at the New York Genome Center giving 125 bp long paired end reads. Samples were sequenced at ~1-5 million reads each. Raw and processed sequencing data are available at GEO under accession number: GSE102998.

### 4C-seq data analysis

All 4C-seq reads were filtered to ensure a match for the bait primers used, then trimmed using a custom script to remove the first 20 nt of each read. Reads were then mapped to the mouse mm9 reference genome using the Burrows-Wheeler aligner (bwa-mem) allowing for up to two mismatches. The package r3Cseq [73] was used to analyze the distribution of signal in *cis*,both near the bait and along chromosome 5. Reads were counted per restriction fragment to obtain the highest possible resolution. Data shown in the main text figures are for the intersection of four male and four female mouse livers, merged according to sex using the intersection option, meaning that the 4C interactions shown are those present in all four samples for a given sex. This produces both a normalized read depth signal (reads per million per restriction fragment) as well as an associated p-value for the interaction taking into account distance from viewpoint and reproducibility across replicates (Figure S7A,C). A comprehensive view of all replicates is presented in Figure S7A. For all pairwise comparisons, a correlation between samples was calculated genome-wide using the UCSC utility bigWigCorrelate with default settings. To account for high signal immediately surrounding the viewpoint, this analysis was only conducted for regions >10 kb from the viewpoint fragment. To analyze more distal *cis* interactions, we first calculated the normalized 4C signal observed per TAD along chromosome 5, in units of RPKM per TAD. We observed a robust logarithmic decay of signal with increasing distance from the viewpoint TAD (Figure 6C; R^2^=0.719). Interacting TADs were defined according to observed over expected 4C signal relative to this background model. Interacting TADs were designated as follows: high, defined as regions with >2-fold enrichment over this background model (observed/ expected); medium, defined as 1.5 to 2-fold enrichment; and low, between 1.5-fold enrichment and 1.5-fold depletion. Non-interacting TADs showed > 1.5-fold depletion of signal. For the *Alb* viewpoint, we identified 17 high, 20 medium, 128 low, and 30 non interacting *cis* TADs along chromosome 5. We required a more comprehensive background model to analyze interactions in *trans*. The tool 4Cker was used for its adaptive windowing and Hidden Markov model approach [71]. The count tables from r3C-seq were merged by sex and imported, then *trans* analysis was conducted with the recommended parameters (k=20). The default output identifies three classes of regions: interacting, low-interacting, and non-interacting. For our analysis, the interacting group was divided into two equal-number groups: high-interacting and medium-interacting, based on 4C interaction strength in the male liver samples. *Trans* interacting regions tend to be large (median size of 1.8 Mb), therefore *trans* interacting TADs were defined as TADs wholly contained within these interacting regions. This corresponds to a total of 659 (high), 618 (medium), 969 (low), and 77 (non) interacting *trans* TADs genome-wide.

### CAC sites and scores

Cohesin-and-CTCF (CAC) sites were defined as CTCF peaks that were present in at least 2 of 4 individual mouse liver samples and that overlapped with a cohesin peak in any liver sample. CAC sites were scanned for a CTCF motif (JASPAR motif MA0139.1) within the CTCF peak coordinates using the FIMO tool in the MEME Suite (version 4.10.0). For a given CAC site, the highest scoring motif occurrence for the canonical core CTCF motif (MA0139.1) was considered. A (+) strand orientation indicates that the motif is found on the (+) genomic strand (Watson strand). Each CAC site was represented by two different scores: a CTCF score = p * (m/10), where p is the CTCF peak strength (MACS2 score) and m is the CTCF motif score, as determined by FIMO; and a cohesin score = p * (m/10), where p is the cohesin (Rad21) peak strength (MACS2 score) and m is the CTCF motif score, as determined by FIMO.

### subTAD prediction method

We applied the published algorithm for CTCF-mediated loop prediction [31] to predict subTAD loop structures. Key modifications to the algorithm include the following: incorporation of cohesin ChIP-seq data in scoring, based on the finding that CTCF signal in the absence of cohesin is not sufficient to predict chromatin interactions [61]; consideration of TAD structure, TSS overlap, and consistency across biological replicates when filtering to obtain the final set of predicted loops; and a final target set of approximately 10,000 subTAD loops, based on experimental results from high resolution Hi-C analyses [12]. First, CAC sites were identified from ChIP-seq data as described above. Next, each chromosome was scanned for putative subTAD loops, formed between a (+) anchor [upstream anchor, i.e., CAC site with a CTCF motif (JASPAR motif MA0139.1) on the (+) strand] at the start of a loop and a (−) anchor [downstream anchor, i.e., CAC site with a CTCF motif on the (−) strand] at the end of a loop, as described for prediction of intra-chromosomal CTCF loops in [31]. Scanning was initiated from the beginning of each chromosome, and a list of putative (+) anchors was generated. Next: (1) if the next CAC site encountered was a (−) anchor, the pair of (+) and (−) anchors was recorded as a putative subTAD loop. The (+) anchor was paired with all subsequent, downstream (−) anchors until another (+) anchor was encountered, at which point the list of putative subTAD loops was closed, ending with the last (−) anchor. Alternatively, (2) if the next CAC sites encountered were (+) anchors, then all such (+) anchors were retained as putative upstream anchors, until the next (−) anchor was reached, and then all such (+) anchors were paired (i.e., assigned to loops) with all of the subsequent, downstream (−) anchors until a new (+) anchor was encountered, as described under (1), at which point the list of putative loops was closed, ending with the last (−) anchor. A new scan for putative subTAD loops was then initiated in a linear fashion, starting from the next (+) anchor until all chromosomes were scanned and a set of putative subTAD loops was obtained. Chromosome scanning for putative subTAD loops was then repeated as described above after removing 10% of the CAC sites - those with the lowest <u>CTCF</u> scores (defined above). Chromosome rescanning was repeated iteratively until the number of putative subTAD loops decreased to as close to 20,000 as possible (removing the lowest scoring loops if needed). A parallel series of iterative scans was carried out, except that 10% of the CAC sites with the lowest <u>cohesin</u> scores (defined above) were removed at each iteration, to generate a second set of ~20,000 putative subTAD loops. The intersection of the two sets of 20,000 putative subTAD loops was then determined. The same iterative process of subTAD loop prediction was carried out independently for each of the n=4 individual mouse livers, based on an analysis of matched CTCF and cohesin (i.e., Rad21) ChIP-seq datasets for each liver. Thus, for each liver sample, a single putative subTAD loop set was generated from the intersection of two sets of predicted CAC-based loops, one using CTCF ChIP-seq strength, and the other using cohesin (Rad21) ChIP-seq strength, with both scores modified by the CTCF motif score. The overlap of these two putative subTAD loop sets was approximately 80%, and ranged from 15,999-16,892 loops for a given liver sample. Additional filters were then applied to remove subTADs that did not contain either a protein-coding TSS or a liver-expressed lncRNA TSS (as defined in [74]), as we were primarily interested in the impact of subTADs on gene expression and regulation. Putative subTAD loops that overlapped >80% of a TAD, or whose (+) and (−) anchors are both TAD anchors (defined below) were also excluded, as these loops could not be distinguished from TAD loops. These two filters further reduced the putative subTAD loop sets to approximately 63% of the original 20,000 (12,395-12,962 loops for a given liver sample). A single merged sample (merged at the fastq file level, separately for CTCF and Rad21) was run through the full pipeline, above, and then sequentially intersected with the set of putative subTAD loops predicted for each individual liver to obtain a final set of 9,543 subTAD loops identified in all 4 livers and also present in the 5^th^ dataset (merged sample). Each subTAD loop was assigned a subTAD loop score equal to the geometric mean of the (+) anchor’s CAC site CTCF score and that of its (−) anchor. A second subTAD loop score, equal to the geometric mean of the (+) anchor’s CAC site cohesin score and that of its (−) anchor, was also assigned. The CTCF and cohesin scores reported for each loop in the final subTAD loop lists (Table S1B) are those obtained from the merged sample.

### RNA-seq Analysis

Gene expression values for liver-expressed protein coding genes are log_2_(FPKM+1) values from [74]. Liver-expressed non-coding genes are expressed in FPKM based on the gene models and expression values from [74]. To express the tissue specificity of a gene’s expression across a panel of 21 mouse tissues (including liver), we used Tau, which was shown to be the most robust in a recent study [46]. Testis was excluded from this analysis because a large proportion of testis-expressed genes are highly tissue specific. For each tissue, the maximum FPKM per gene between the two replicates was used. These FPKM values were log transformed and a Tau value, ranging from 0 to 1, was calculated, where 1 represents high tissue specificity: T = Σn_i_ = 1(1−y_i_)n−1, where y_i_ = x_i_max_1≤i≤n_(x_i_), n is the number of tissues, and x_i_ is the expression of the gene in tissue i.

### General Hi-C Processing

Hi-C data was processed using the HiC-Pro package (version 2.7.0) [75] for mapping and read filtering, followed by Homer (version 4.8) for downstream analyses such as PCA analysis and aggregate contact profiles. Biological replicates were merged to increase read depth. The default Homer background model was used for all datasets, where the expected frequency of interactions takes into account read depth between interacting bins and genomic distance. PCA was conducted using Homer with the command “runHiCpca.pl -res 10000 -cpu 4-genome mm9” to generate genome wide eigenvalues at 10 kb resolution. The values changed marginally at 20, 40, or 50 kb, but the sign of the eigenvalue was unaffected, i.e., there was no impact on whether a TAD was assigned as A compartment or B compartment.

### Peak distribution within TADs

Published TAD coordinates in mouse liver [3] were converted from mouse genome mm10 to mm9 using liftover with default parameters. Each TAD was then divided into 100 equal-sized bins using the Bedtools command makewindows. Next, these bins were compared to the peak positions of various publicly available ChlP-seq datasets using the Bedtools coverage command, and the number of peaks per bin was counted. This resulted in a string of 100 values for each TAD, representing the number of ChlP-seq peaks per bin, where the first value is the start of the TAD and the last value is the end of the TAD. Conducting this analysis across all TADs yielded a matrix of 3,617 rows (one per TAD) × 100 columns (one per bin). To generate the aggregate profiles shown in Figure 1A-1D, the sum of each column was taken and then normalized to account for differences in total peak count for the different samples, factors, and chromatin marks analyzed. Normalization was conducted by taking the average of the center five bins (bins 48-52) and dividing the bin sums by this normalizing factor. This allows the y axis to represent bin enrichment relative to the center of the TAD, as shown.

### TAD activity and compartment assignment

‘TAD boundaries’ were defined at single nucleotide resolution as the end of one TAD and the start of another (as defined in [3]), thus excluding the start of the first TAD in each chromosome and the end of the last TAD. In contrast, all references to ‘TAD anchors’ refers to the CTCF sites most likely to be anchoring TAD loops based on distance from the boundary and proper orientation (as described in TAD Anchor Identification). Data sources for all ChIP-seq, GRO-seq, Hi-C, and other datasets are described in Table S4. H3K9me3, H3K27me3, H2AK5ac, and H3K36me3 marks were processed from the raw sequencing data (fastq files) through the standard ChIP-seq pipeline, above. H3K9me3 and H3K27me3 mark data were expressed as log_2_(ChIP/IgG signal). Lamina-associated domain coordinates and GRO-seq data were downloaded as pre-processed data. Heat maps were generated using Deeptools reference point, with a bin size of 10 kb. TAD boundaries were grouped according to k-means clustering (k=4) using signal within a 1 Mb window from 3 datasets: H3K9me3, H2AK5ac, and the eigenvalue of the Hi-C PCA analysis (above). Based on these clusters, TADs were classified as active, weak active, weak inactive, or inactive, as follows. If both the start and end boundary of a given TAD were classified as active, then the TAD was designated active. Specifically, a TAD was considered “active” if the boundary at the start of a TAD fell into clusters 1 or 2 (as marked in Figure 1E) and the boundary at the end of the same TAD fell into clusters 1 or 3. The corresponding metric was applied to identify inactive TADs. If the two ends of a given TAD were not in agreement, then the TAD was designated weakly active if the median Hi-C PC1 eigenvalue within the TAD was positive, or weakly inactive if the median Hi-C PC1 eigenvalue was negative. Gene expression and tissue specificity metrics represent expression or Tau values of genes whose TSS overlap active or inactive TADs.

### Additional Hi-C analysis

Contact profiles around TAD, subTAD, and non-loop-anchor CTCF sites were generated using Homer (v4.9) using the command analyzeHiC and the options “-size 500000 -hist 5000” to generate interaction profiles for 1 Mb windows around CTCF sites with 5 kb resolution. TAD and subTAD anchors were split into left and right anchors if they were at the start or the end of the predicted loop. Nonanchor CTCF sites were defined as any remaining CTCF sites from the merged sample that also contained a CTCF motif. Left and right groupings were determined based on the orientation of the strongest CTCF motif within the non-anchor peak regions. The inward bias index (IBI) was modified from the more genome-wide directionality index (DI) described in [1]. Both DI and IBI use a chi-squared statistic to determine the extent to which Hi-C reads from a given region have a strong upstream or strong downstream bias. While DI is genome wide, IBI focuses on the directionality of *cis* interactions (within 2 Mb) from a 25 kb window immediately downstream of a CTCF peak relative to the motif orientation. A large positive value indicates a strong interaction bias towards the loop center, as the motif orientation would predict. Values close to zero indicate a roughly equal distribution of interactions upstream and downstream. By orienting the sign of the IBI value relative to the CTCF motif directionality, we were able to group left and right loop anchors together.

### TAD anchor identification

TAD anchors were predicted using a similar approach. The merged list of CTCF peaks was filtered to only consider peaks that were found across all four biological replicates, that contained CTCF motifs, and that were within 50 kb of a TAD boundary, as defined previously for mouse liver [3]. This 50 kb distance was chosen based on the ambiguity of binned Hi-C data to more accurately determine the precise TAD boundary. Then, for each TAD boundary, all pairs of (+) and (−) CTCF peaks were considered and scored based on their combined distance to the called TAD boundary. Pairs of “+/−” peaks that were comprised of a (+) anchor upstream of a (−) anchor (i.e., CTCF peak pairs that were not divergently oriented) were considered an invalid combination to define the end of one TAD and the beginning of the next TAD, and were not considered. The valid pairs with the shortest combined distance to the previously defined liver TAD boundary [3] were retained and all others were removed. If no valid pair for a TAD boundary was identified, the single CTCF peak closest to the TSD boundary was retained as the TAD anchor. A complete listing of TAD anchors can be found in Table S1, and a listing of inter-TAD regions and associated gene ontology analysis is presented in Table S3.

### Alternative loop anchor analysis

We sought to compare the relative insulation of loops identified by our computational approach to alternative loops identified using the original core algorithm [31]. This provides an objective measure to compare the performance of each implementation in identifying TAD-like loops and loop anchors within TADs. To this end, we used the complete CTCF peak list from the merged sample as input and implemented the loop prediction algorithm exactly as described previously (60% proportional peak cutoff, CTCF signal + motif scores as above, retaining only loops <200 kb) [31]. As summarized in Figure S2B, this analysis yielded many more loops (60,678), which were shorter (median size of 61 kb) and showed less overlap with cohesin-mediated loops in the mESC ChIA-PET dataset (25.5% versus 68.3% for subTADs). 59% of the subTADs characterized in our study are found in this larger loop list (termed “60k loops”). To characterize loops unique to the 60k loop set, we had to first filter out anchors found subTAD or TAD loops (to insure that each list was mutually exclusive, as above). Any 60k loop anchor within 50 kb of a TAD boundary was excluded from downstream analysis. Similarly, we also excluded any 60k loop with at least one anchor that coincided with a subTAD anchor. These mutually exclusive lists of subTAD anchors and the filtered set of 60k loop anchors (25,983 loop anchors in total; representing a subset of “Non Anchor CTCF” in the main text) were then compared based on insulation of repressive histone marks (see “Repressive histone mark insulation”, below) and Hi-C interaction profiles (see “Additional Hi-C analysis”, above). Figure S2B,C presents these insular features of subTAD anchors compared to alternative loop anchors that are not subTAD or possible TAD anchors.

### Anchor/Loop overlap

CTCF ChIP-seq data for 15 non-liver tissues from the ENCODE Project were downloaded (https://genome.ucsc.edu/cgi-bin/hgTrackUi?db=mm9&g=wgEncodeLicrTfbs) and intersected with replicates to form a single peak list for each tissue [42]. These single peak lists per tissue were then compared to liver CTCF peaks using the Bedtools multiinter command with the -cluster option to generate a union CTCF peak list for all tissues with a score representing the number of tissues in which a peak is present. “Lone” CTCF (CTCF sites lacking cohesin bound), other/non-anchor CAC sites, TAD anchors, and subTAD anchors were compared to this list to generate the histograms in Figure 2C (See also, Table S1C). Knockdown-resistant cohesin binding sites in liver were defined as Rad21 ChIP-seq peaks found in both wild-type (WT) and Rad21^+/−^ mouse liver, with knockdown-sensitive sites defined as Rad21 peaks found in WT liver that are absent in Rad21^+/−^ liver [17]. Similarly, knockdown-resistant cohesin binding sites in MEFs were defined as Smc1a ChIP-seq peaks present in both WT [16] and Stag1-knockout MEFs [76]. Knockdown-sensitive sites were defined as Smc1a peaks found in WT MEFs that are absent in Stag1-knockout MEFs. Phastcons 30-way vertebrate conservation scores were downloaded from the UCSC table browser and converted to BigWig tracks using ucscutils (version 20130327; ftp://hgdownload.cse.ucsc.edu/goldenPath/mm9/phastCons30way/vertebrate). Comparisons to mESC Smc1a ChIA-PET and Smc1a Hi-ChIP datasets [5, 50] were based on merged replicates, and reciprocal overlaps with subTADs were required (Bedtools intersect -wa -u -r -f 0.8 -a subTAD.bed -b mESC.bed). mESC interactions were filtered to only include anchor regions that also contain a CTCF peak, to exclude CNC-mediated enhancer-promoter interactions.

### Repressive histone mark insulation

To determine if a TAD or subTAD anchor CAC showed more insulation, or less insulation, than other classes of CTCF or cohesin binding sites, we used Jensen Shannon Divergence (JSD; [51]) to quantify the insulation of H3K27me3 and H3K9me3 ChIP-seq signals. Specifically, regions 10 kb upstream and 10 kb downstream of each peak in a given peak list (i.e., TAD anchors, CNC, etc.) were each divided into 50 bins of 200 bp each. The number of H3K27me3, H3K9me3, or IgG ChIP-seq reads within each bin was tallied, resulting in a vector of 50+50 values for each peak region. These were then compared to two test vectors representing complete (maximal) insulation: fifty 0’s followed by fifty 1’s, and fifty 1’s followed by fifty 0’s. These are theoretical representations of low signal upstream of the peak followed by high signal downstream, and vice versa. Using a custom python script, the similarity between the experimentally derived vector and each of the test vectors was calculated, where a lower value represents less divergence from the test vector. The cumulative frequency distribution per group (anchors, CAC, CNC, etc.) is presented for the most similar test vector per peak in Figure 3D, 3E (K27me3 and K9me3) and Figure S4F (IgG). Heat maps show ChIP signal Z-transformed data across all CTCF-bound regions.

### Five Class DHS model

The ~70,000 open chromatin regions (DHS) previously identified in mouse liver [54] were classified based on ChIp-seq signals for H3K4me1, H3K4me3, and CTCF within 1 kb of each DHS summit, obtained using the refinepeak option in MACS2. The general schematic is shown in Figure S5A. Promoter DHS were defined as DHS with a > 1.5-fold ratio of H3K4me3 relative to H3K4me1 ChIP signal; enhancer DHS were defined as DHS with a <0.67-fold ratio of H3K4me3 relative to H3K4me1 signal. Both DHS sets were filtered to remove DHS with <4 reads per million for each mark after subtracting IgG signal (Figure 4A). These cutoff values leave two remaining DHS groups, one with a roughly equal ratio between the two histone-H3 marks, and one with low signal (<4 reads per million) for both marks. The former DHS were classified as weak promoter DHS, based on their close proximity to TSS and the low expression of neighboring genes (Figure S5B, S5C). The remaining DHS group, characterized by low ChIP-seq signals, was largely intergenic but showed weak to undetectable levels of canonical histone marks. Low signal regions that overlapped a CTCF site with higher CTCF ChIP-seq signals than H3K4me1 signals were classified as potential insulators (Figure S5A). The remaining regions were designated weak enhancer-DHS based on their distance from TSS and their low levels of H3K27ac compared to the enhancer-DHS group (Figure S5B). The majority of promoter-DHS and weak promoter-DHS were <1 kb from a TSS (Figure S5B). To compare the level of expression for genes with promoter-DHS or weak promoter-DHS, the TSS was required to be within 10 kb of the DHS summit (Figure S5C). Any gene with both a weak promoter-DHS and a promoter-DHS within 10 kb was categorized as being regulated by a promoter-DHS; thus, there was no overlap between weak promoter-DHS-regulated genes and promoter-DHS-regulated genes.

### Comparison of DHS classes across tissues

All available mouse tissue DNase-seq peak regions were downloaded from the ENCODE Project website (https://www.encodeproject.org/; [42]), blacklist regions were removed, and the lists were merged to form a single reproducible peak list for each tissue, as follows. Due to variable replicate numbers across tissues, the following cutoffs were used to form merged DHS lists for each tissue. If a tissue had only two replicates (as was the case for 12 of the 20 non-liver tissues), we required that the DHS be present in both replicates. If a tissue had 3 or 4 replicates, then the DHS were required to be present in all or all but one replicate (this was the case for 7 of the 20 non-liver tissues). For whole brain tissue, the merged peak list required that a DHS was present in at least 5 of the 7 replicates. These regions were compared to each other using the Bedtools multiinter command with the -cluster option to generate a union DHS peak list for all tissues, where the score column represents the number of non-liver tissues in which a given region was found. For all liver DHS assigned to one of the above five DHS classes (Table S2A), each liver DHS summit was mapped to this all tissue union peak list, allowing only one match per summit up to 150 nt away. If a given liver DHS summit was >150 nt from the nearest DHS in any other tissue, it was given a score of “0” for liver-specificity. Otherwise the score represents the number of mouse tissues that the closest DHS was found in.

### Super-enhancer identification

Super-enhancers were identified using the ROSE (Ranked Order of Super Enhancers) software package (http://younglab.wi.mit.edu/super_enhancer_code.html). ROSE takes a list of enhancer regions and mapped read positions as input to identify highly active clusters of enhancers. Default options were used, including 12.5 kb as the maximum distance for grouping (stitching) enhancers into putative super-enhancers, as well as reads per million normalization for all H3K27ac ChIP signal used for ranking enhancer clusters. The set of all enhancer-DHS and weak enhancer-DHS regions from the five class DHS model described above (Table S2A) was used as the region input list. A set of 19 publicly available H3K27ac ChIP-seq datasets from mouse liver was used as signal input (see Table S4 for sample information). This set includes male, female, and circadian time course data [53]. A strict intersection of super-enhancers identified across all 19 samples was used to define a set of 503 “core” super-enhancers in mouse liver using the Bedtools multiintersect command, as shown in Figure S5E. Any enhancer cluster (i.e., constituent enhancers within 12.5 kb, as above) not identified as a super-enhancer in any sample was termed a typical enhancer.

Gene targets for enhancers were assigned as the nearest gene up to a maximum distance cutoff of 25 kb. Gene expression values and tissue specificity were defined as described above. Aggregate plots were generated using Deeptools (version 2.3.3). In Figure 4F, the scale-regions option of Deeptools was used to scale superenhancers and typical enhancers to their median sizes of 44 kb and 1 kb, respectively. Figure S5F used the reference-point mode of Deeptools and shows GRO-seq signal that overlaps eRNA loci 984 as defined previously [58]. Super-enhancer and typical enhancer coordinates for mESC and ProB cells are from [57].

## DATA AVAILABILITY

Data generated and used in this study has been deposited in the Gene Expression Omnibus (GEO) under accession number GSE102999 (https://www.ncbi.nlm.nih.gov/geo/query/acc.cgi?acc=GSE102999). ChIP-seq data are available under the subseries GSE102997(https://www.ncbi.nlm.nih.gov/geo/query/acc.cgi?acc=GSE102997). 4C-seq data are available under the subseries GSE102998 (https://www.ncbi.nlm.nih.gov/geo/query/acc.cgi?acc=GSE102998). Other datasets used in this study are listed in Table S4.

## SUPPLEMENTARY DATA

Supplementary Data are available online.

## ACKNOWLEDGEMENT

We thank Andy Rampersaud and Tisha Melia for their work building and standardizing the ChIP-seq and RNA-seq pipelines used in this study. We also thank Aram Shin for first piloting CTCF and Rad21 ChIP-seq experiments in this lab.

## Supplemental Information

### Figure Outline—

**Figure S1:**
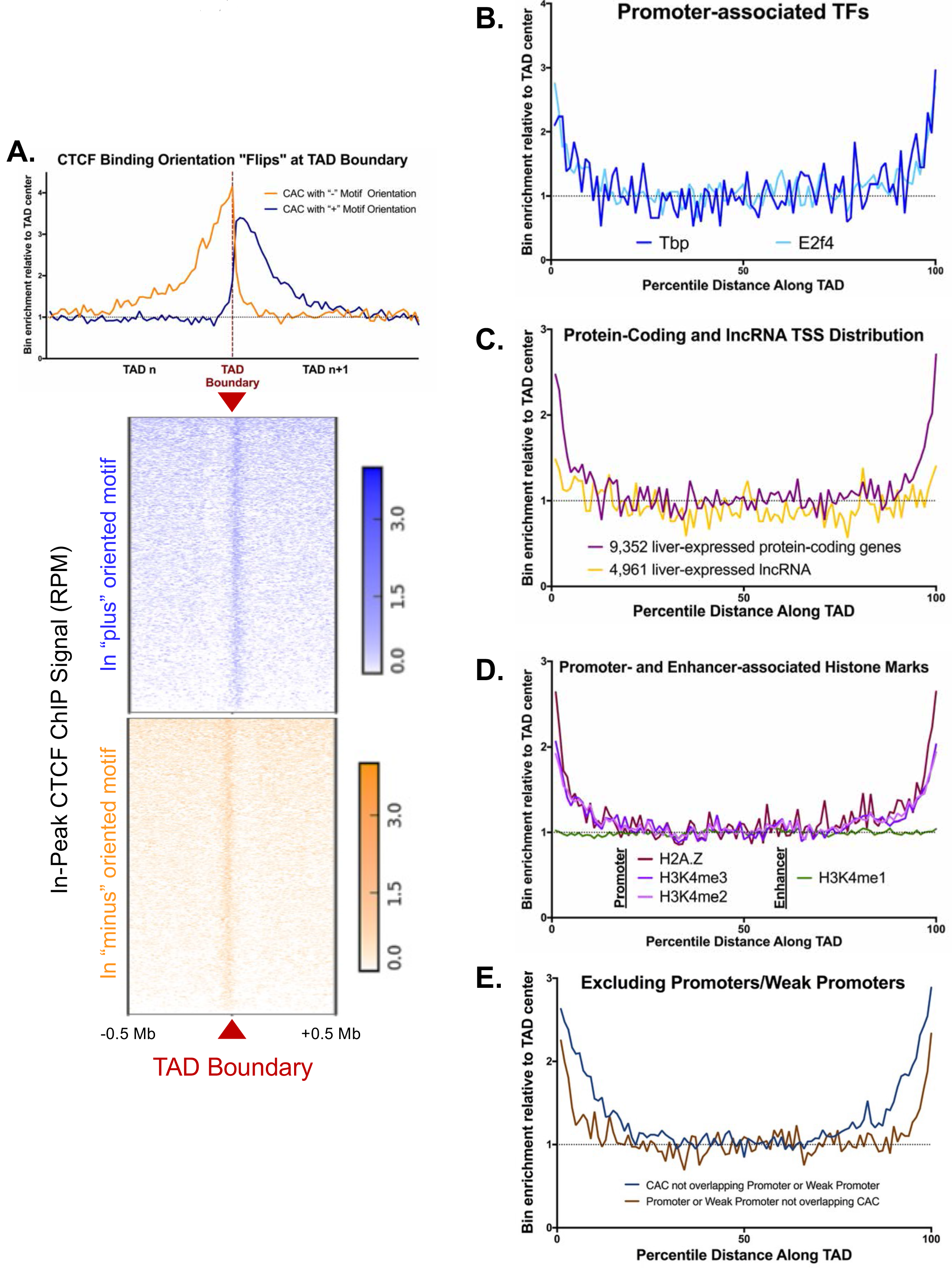
Additional Features of TAD boundaries

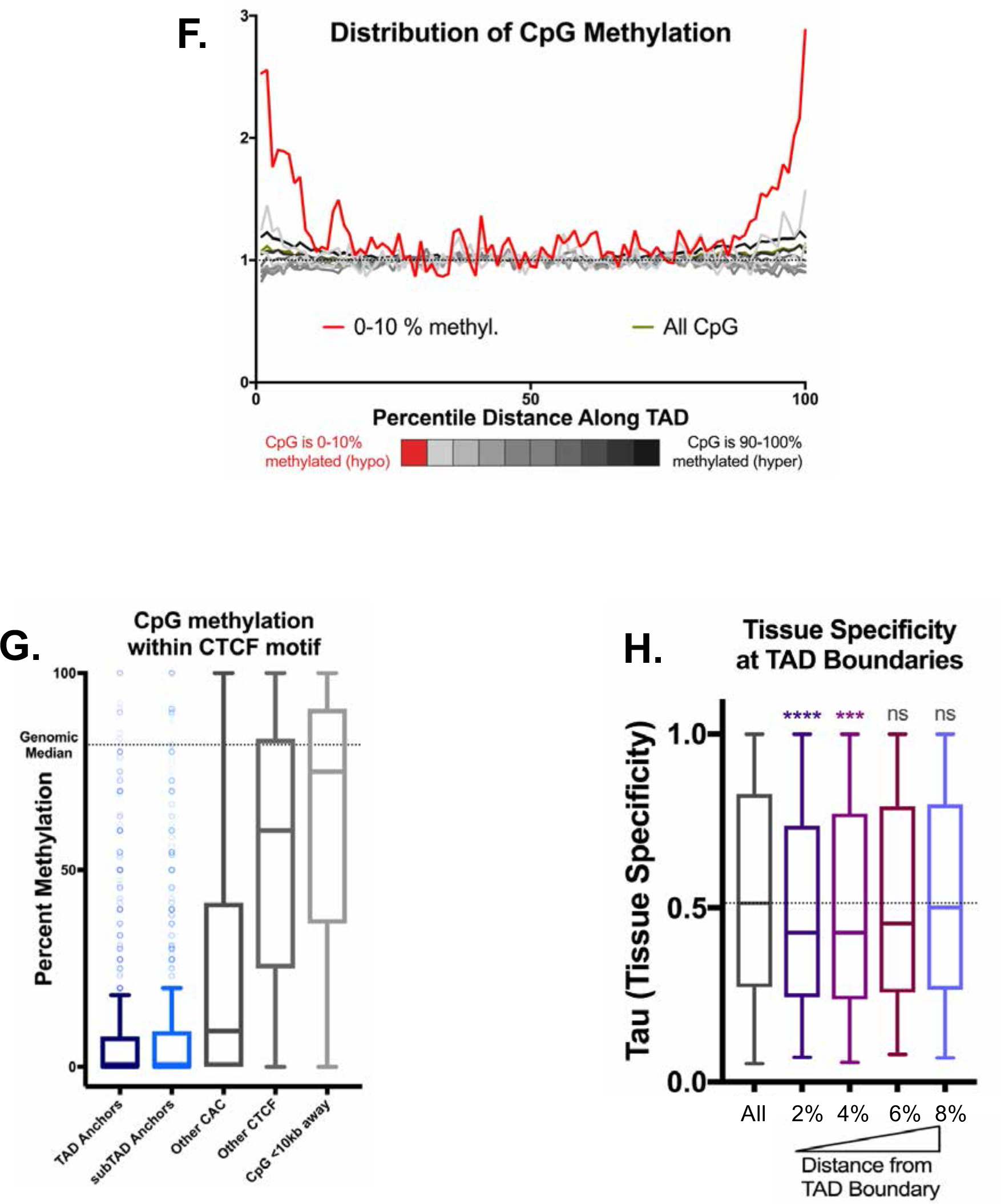

**Figure S2:**
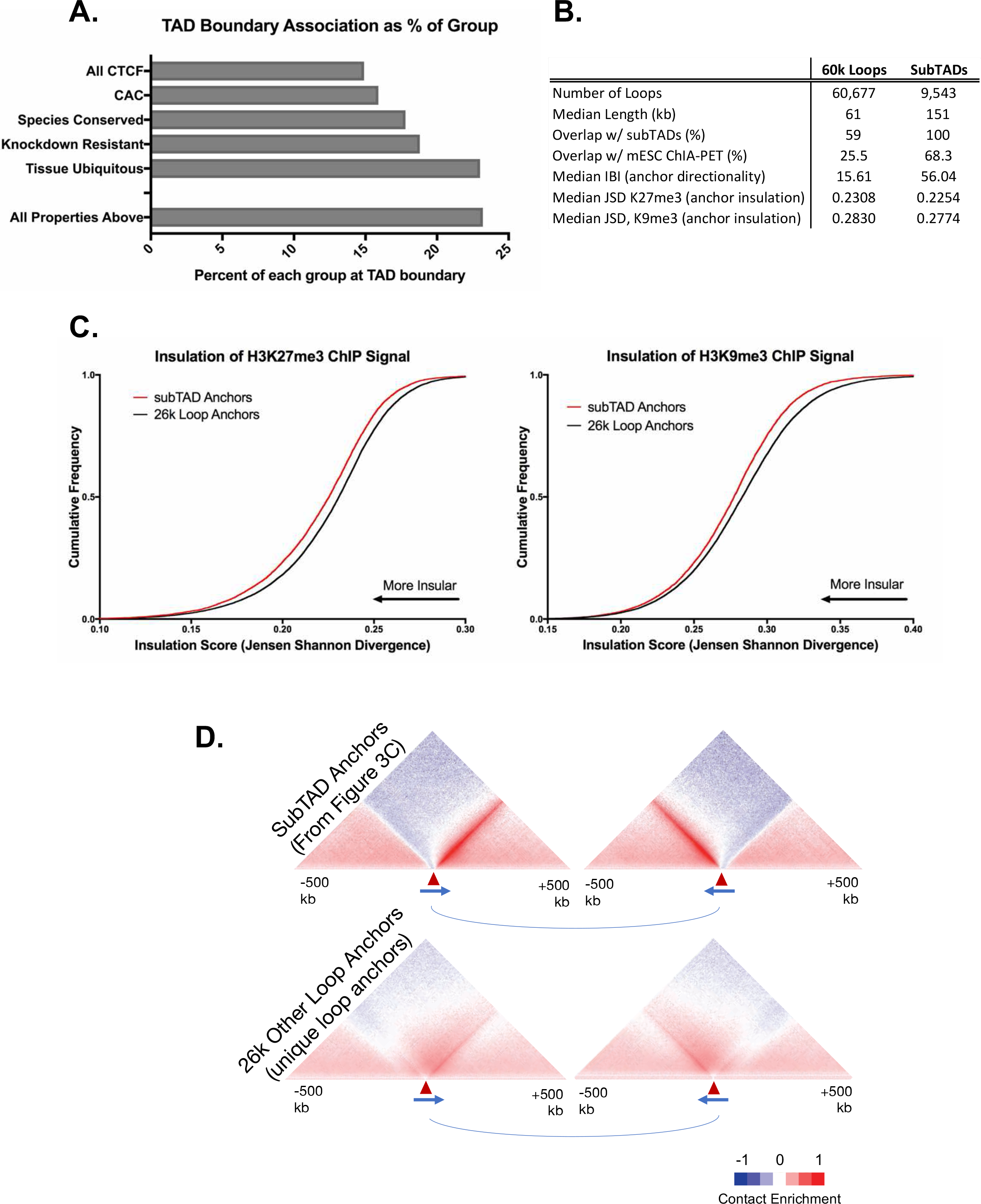
Comparison to prior CTCF loop prediction approaches

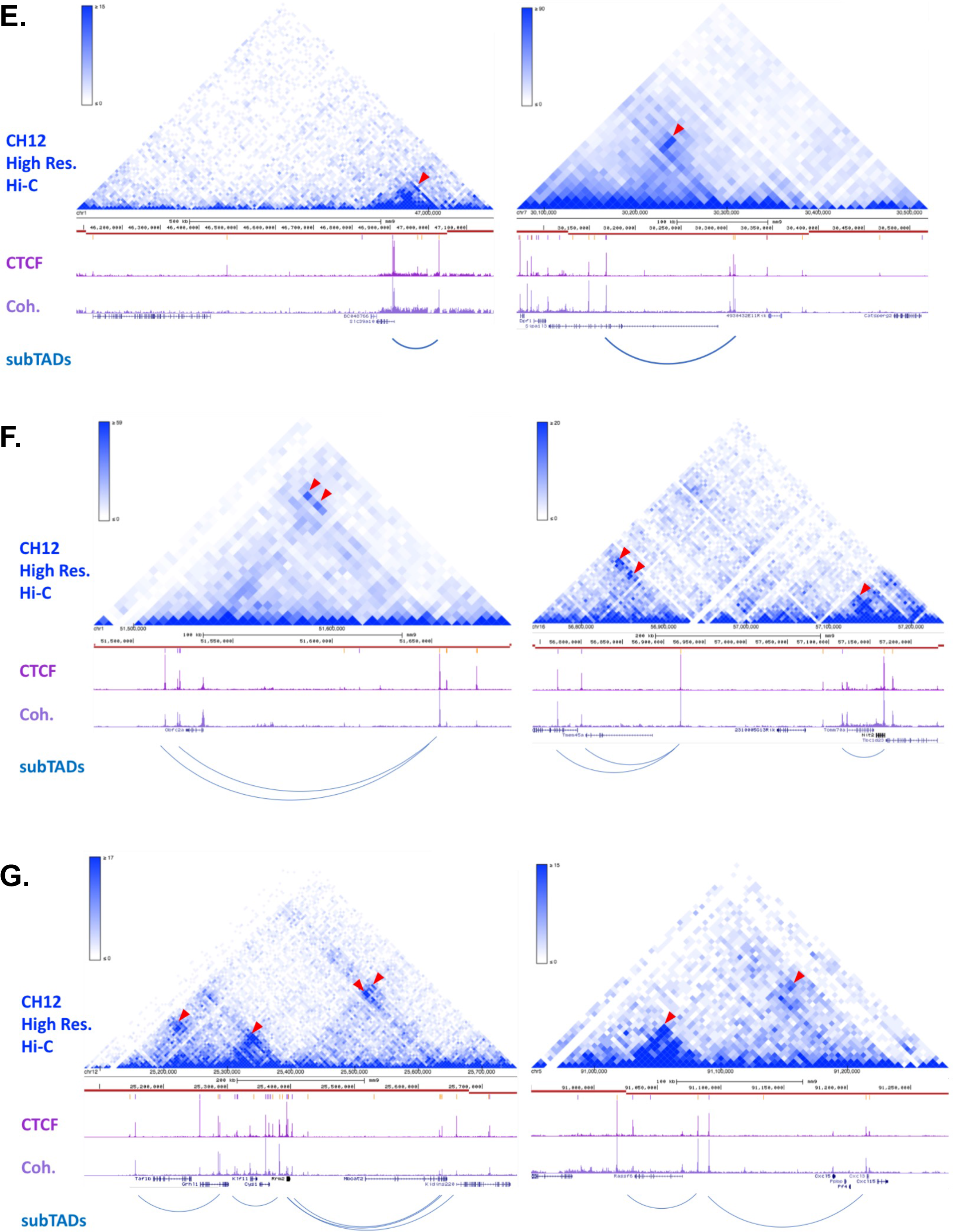

**Figure S3:**
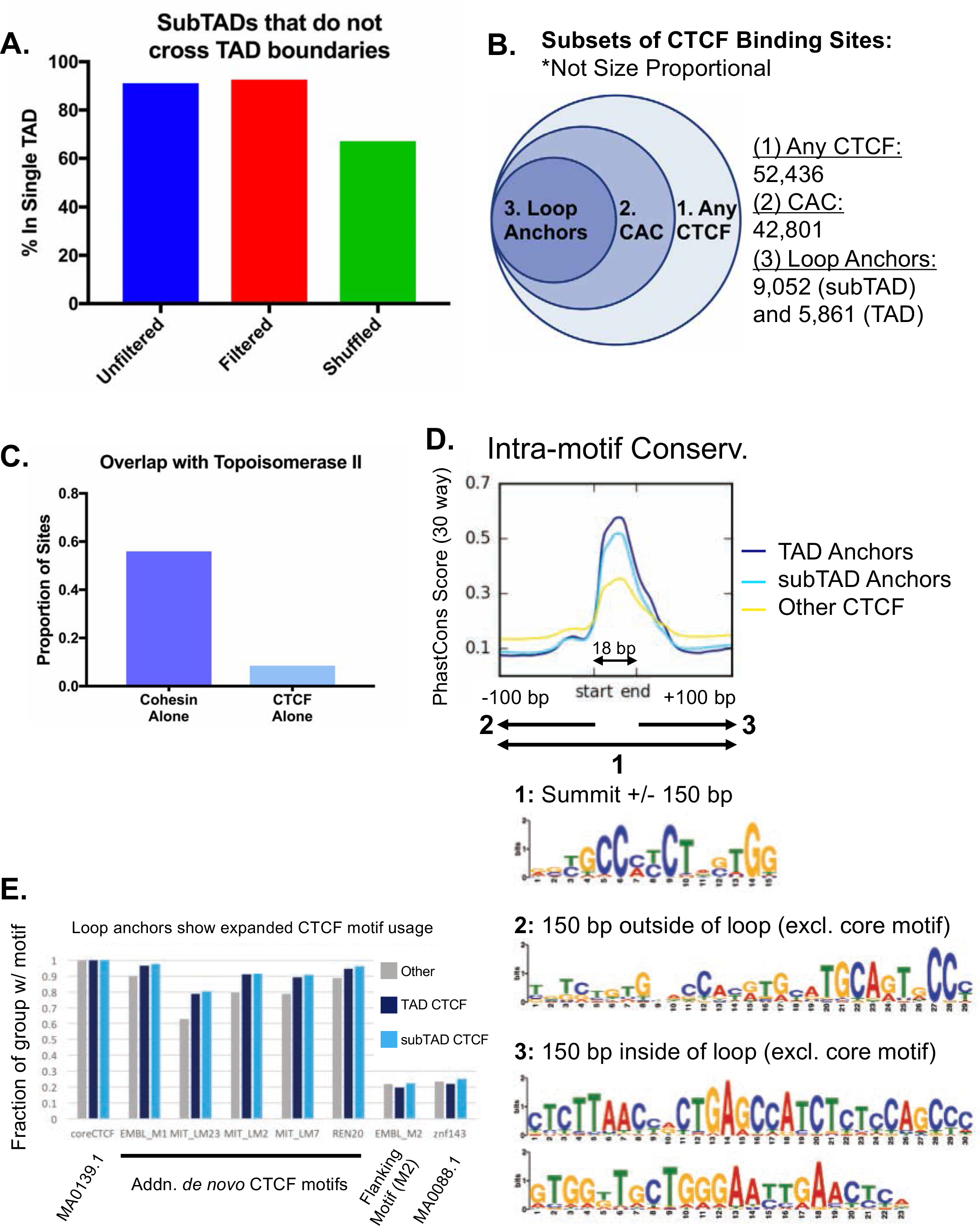
Subclasses of CTCF binding events in relation to predicted loops

**Figure S4:**
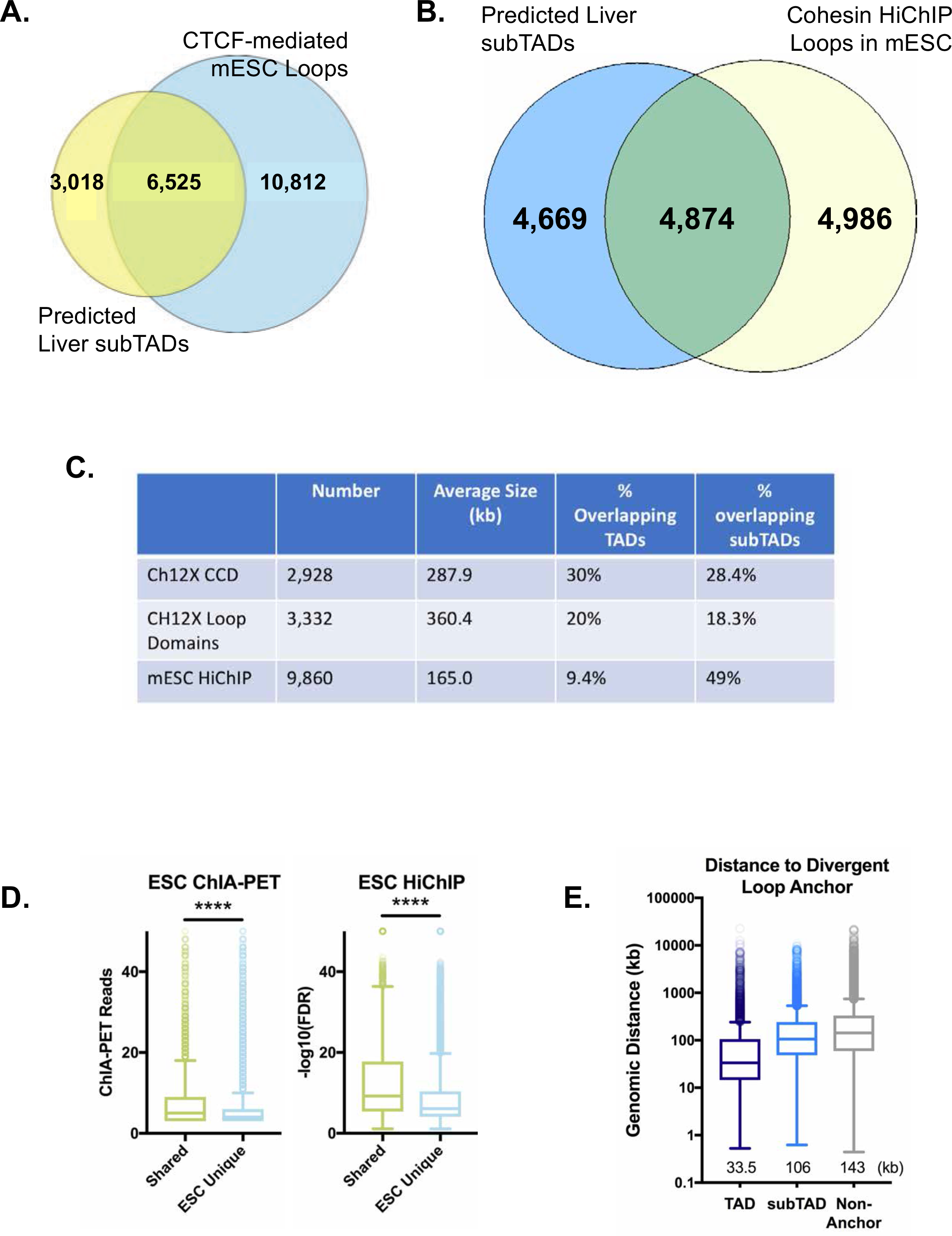
Additional features of subTADs and their insulation

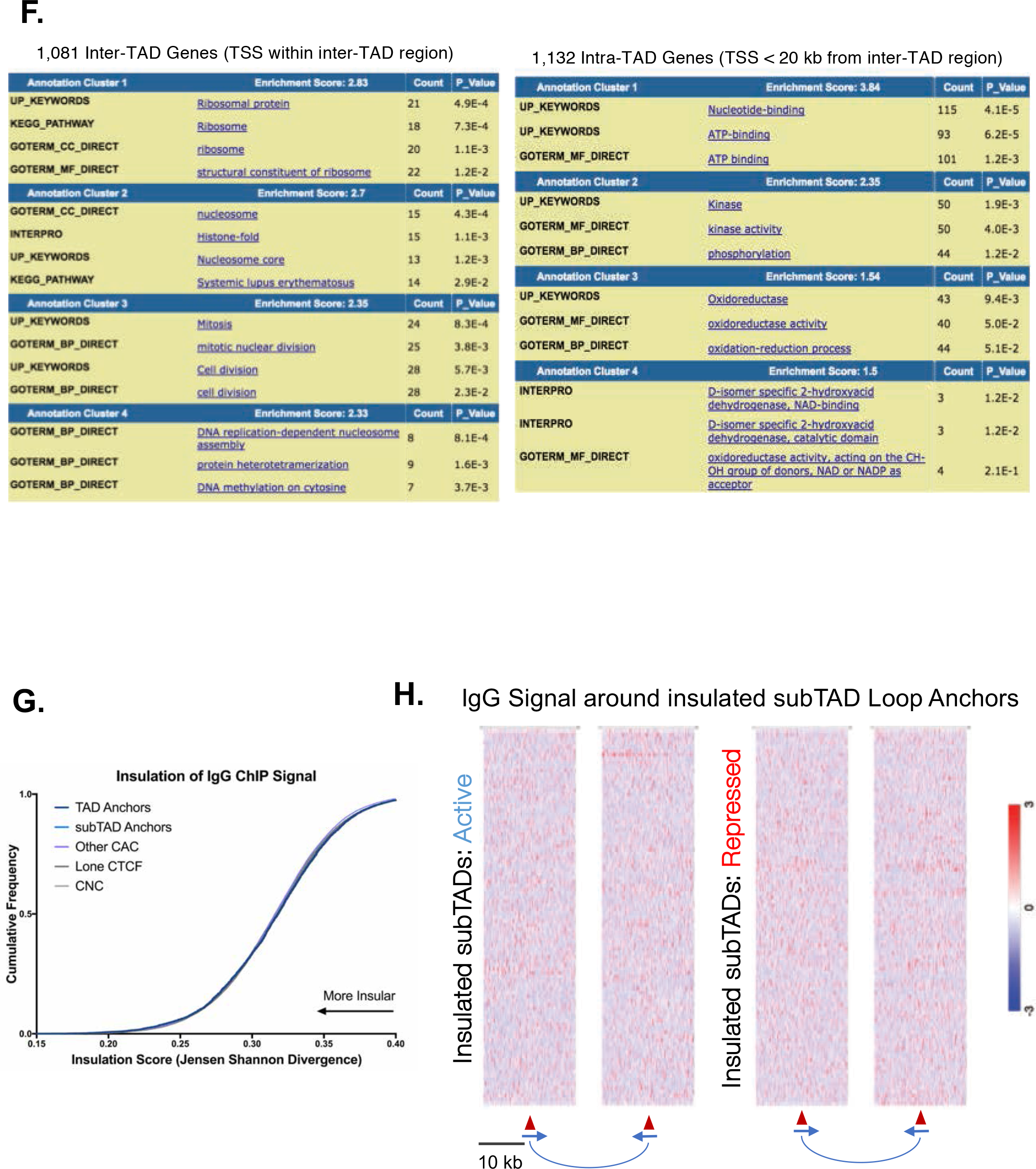

**Figure S5:**
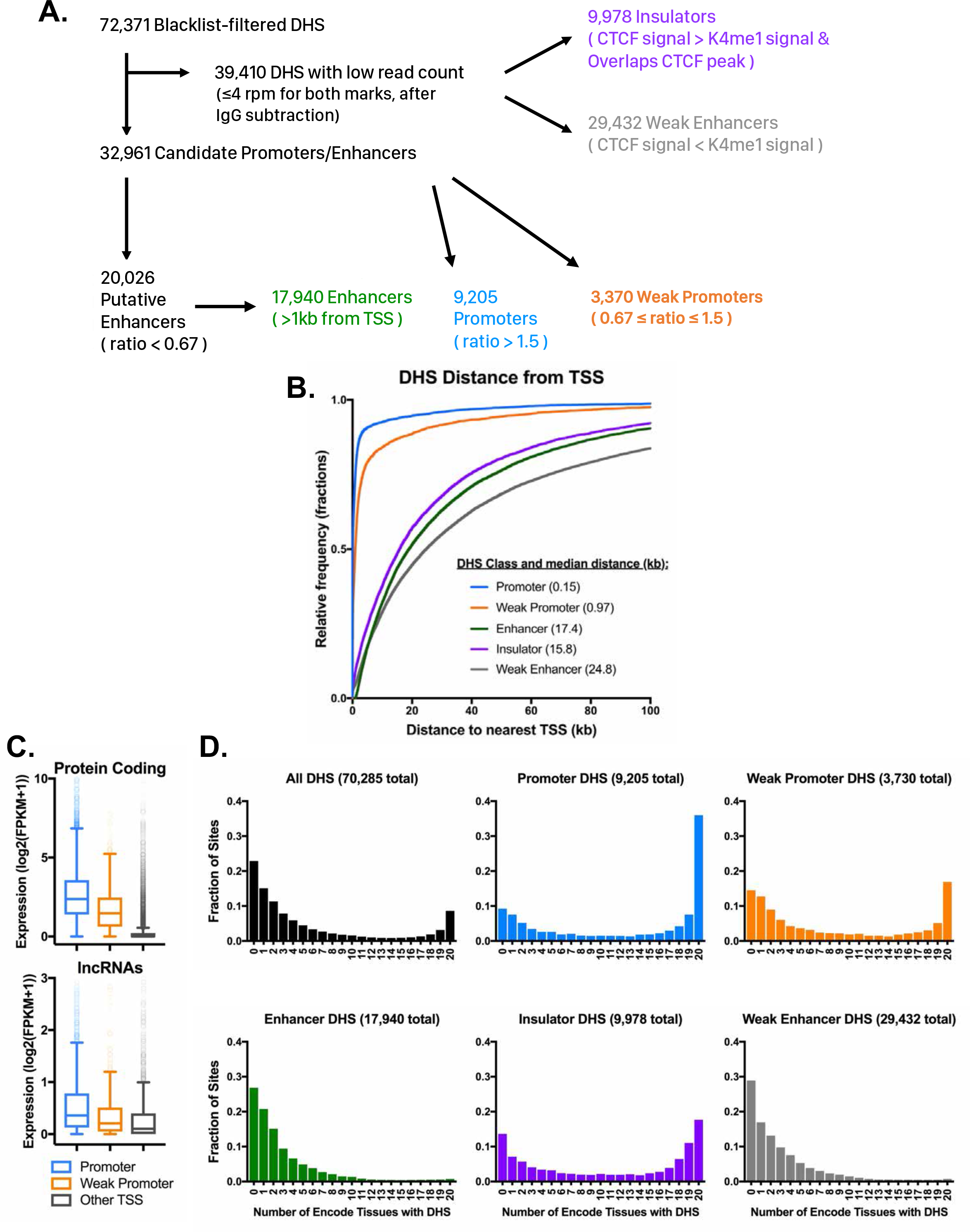
Characteristics of the 5 classes of DHS in mouse liver and

**Figure S6:**
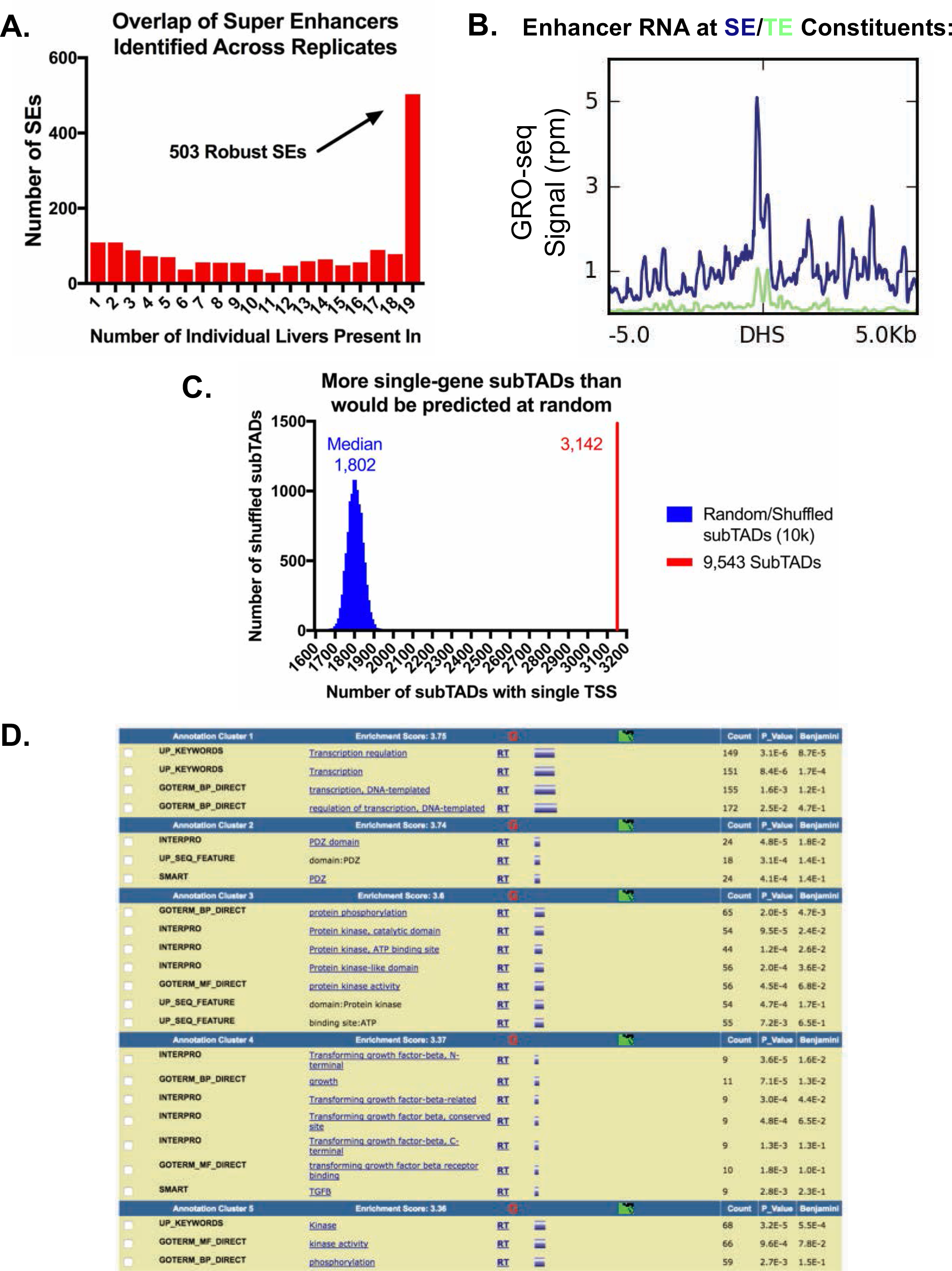
SE single-TSS subTAD features

**Figure S7:**
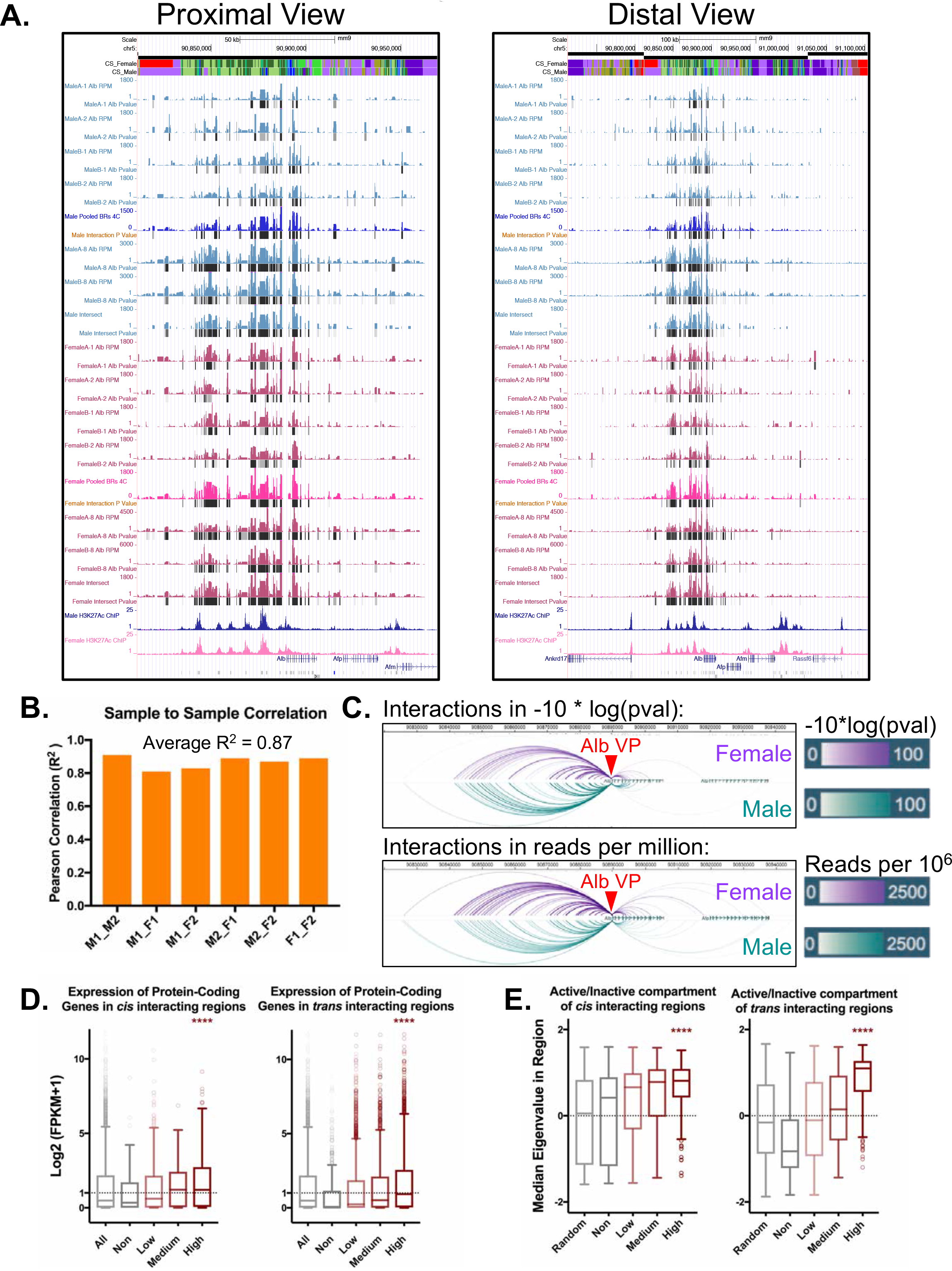
Alb 4C-seq replicates and cis/trans signal distribution

### Extended Methods—

#### Table Outline—

Table S1: TAD coordinates with A/B compartment assignment, subTAD coordinates with loop score, CTCF peak list with CAC classification, CNC peak list, 939 liver-expressed genes in inactive TADs

Table S2: DHS classification, Core Super Enhancers, GO Terms for superenhancer genes, GO terms for typical-enhancer genes

Table S3: Inter-TAD regions/genes, single TSS subTADs, and GO term enrichment

Table S4: Data Sources

## REFERENCES

1. Dixon JR, Selvaraj S, Yue F, Kim A, Li Y, Shen Y, Hu M, Liu JS, Ren B: Topological domains in mammalian genomes identified by analysis of chromatin interactions. Nature 2012, 485:376–380.

2. Nora EP, Lajoie BR, Schulz EG, Giorgetti L, Okamoto I, Servant N, Piolot T, van Berkum NL, Meisig J, Sedat J,et al: Spatial partitioning of the regulatory landscape of the X-inactivation centre. Nature 2012, 485:381–385.

3. Vietri Rudan M, Barrington C, Henderson S, Ernst C, Odom DT, Tanay A, Hadjur S: Comparative Hi-C reveals that CTCF underlies evolution of chromosomal domain architecture. Cell Rep 2015, 10:1297–1309.

4. Dixon JR, Jung I, Selvaraj S, Shen Y, Antosiewicz-Bourget JE, Lee AY, Ye Z, Kim A, Rajagopal N, Xie W,et al: Chromatin architecture reorganization during stem cell differentiation. Nature 2015, 518:331–336.

5. Dowen JM, Fan ZP, Hnisz D, Ren G, Abraham BJ, Zhang LN, Weintraub AS, Schuijers J, Lee TI, Zhao K, Young RA: Control of cell identity genes occurs in insulated neighborhoods in mammalian chromosomes. Cell 2014, 159:374–387.

6. Sexton T, Schober H, Fraser P, Gasser SM: Gene regulation through nuclear organization. Nat Struct Mol Biol 2007, 14:1049–1055.

7. Lieberman-Aiden E, van Berkum NL, Williams L, Imakaev M, Ragoczy T, Telling A, Amit I, Lajoie BR, Sabo PJ, Dorschner MO,et al: Comprehensive mapping of long-range interactions reveals folding principles of the human genome. Science 2009, 326:289–293.

8. Pope BD, Ryba T, Dileep V, Yue F, Wu W, Denas O, Vera DL, Wang Y, Hansen RS, Canfield TK,et al: Topologically associating domains are stable units of replication-timing regulation.Nature 2014, 515:402–405.

9. Nora EP, Dekker J, Heard E: Segmental folding of chromosomes: a basis for structural and regulatory chromosomal neighborhoods? Bioessays 2013, 35:818–828.

10. Le Dily F, Bau D, Pohl A, Vicent GP, Serra F, Soronellas D, Castellano G, Wright RH, Ballare C, Filion G,et al: Distinct structural transitions of chromatin topological domains correlate with coordinated hormone-induced gene regulation. Genes Dev 2014, 28:2151–2162.

11. Fudenberg G, Imakaev M, Lu C, Goloborodko A, Abdennur N, Mirny LA: Formation of Chromosomal Domains by Loop Extrusion. Cell Rep 2016, 15:2038–2049.

12. Rao SS, Huntley MH, Durand NC, Stamenova EK, Bochkov ID, Robinson JT, Sanborn AL, Machol I, Omer AD, Lander ES, Aiden EL: A 3D map of the human genome at kilobase resolution reveals principles of chromatin looping. Cell 2014, 159:1665–1680.

13. Sanborn AL, Rao SS, Huang SC, Durand NC, Huntley MH, Jewett AI, Bochkov ID, Chinnappan D, Cutkosky A, Li J,et al: Chromatin extrusion explains key features of loop and domain formation in wild-type and engineered genomes. Proc Natl Acad Sci U S A 2015, 112:E6456–6465.

14. Phillips JE, Corces VG: CTCF: master weaver of the genome. Cell 2009, 137:1194–1211.

15. Zuin J, Dixon JR, van der Reijden MI, Ye Z, Kolovos P, Brouwer RW, van de Corput MP, van de Werken HJ, Knoch TA, van IWF,et al: Cohesin and CTCF differentially affect chromatin architecture and gene expression in human cells. Proc Natl Acad Sci U S A 2014, 111:996–1001.

16. Kagey MH, Newman JJ, Bilodeau S, Zhan Y, Orlando DA, van Berkum NL, Ebmeier CC, Goossens J, Rahl PB, Levine SS,et al: Mediator and cohesin connect gene expression and chromatin architecture. Nature 2010, 467:430–435.

17. Faure AJ, Schmidt D, Watt S, Schwalie PC, Wilson MD, Xu H, Ramsay RG, Odom DT, Flicek P: Cohesin regulates tissue-specific expression by stabilizing highly occupied cis-regulatory modules. Genome Res 2012, 22:2163–2175.

18. Uuskula-Reimand L, Hou H, Samavarchi-Tehrani P, Rudan MV, Liang M, Medina-Rivera A, Mohammed H, Schmidt D, Schwalie P, Young EJ,et al: Topoisomerase II beta interacts with cohesin and CTCF at topological domain borders. Genome Biol 2016, 17:182.

19. Ong CT, Corces VG: CTCF: an architectural protein bridging genome topology and function. Nat Rev Genet 2014, 15:234–246.

20. Guo Y, Xu Q, Canzio D, Shou J, Li J, Gorkin DU, Jung I, Wu H, Zhai Y, Tang Y,et al: CRISPR Inversion of CTCF Sites Alters Genome Topology and Enhancer/Promoter Function. Cell 2015, 162:900–910.

21. Heath HRdA C; Sleutels, F; Dingjan, G; van de Nobelen S; Jonkers, I; Ling, K; Gribnau, J; Renkawitz, R; Grosveld, F; Hendriks, R. W.; Galjart, N: CTCF regulates cell cycle progression of aB T cells in the thymus. EMBO J 2008, 27:2839–2850.

22. White JK, Gerdin AK, Karp NA, Ryder E, Buljan M, Bussell JN, Salisbury J, Clare S, Ingham NJ, Podrini C,et al: Genome-wide generation and systematic phenotyping of knockout mice reveals new roles for many genes. Cell 2013, 154:452–464.

23. Xu H, Balakrishnan K, Malaterre J, Beasley M, Yan Y, Essers J, Appeldoorn E, Tomaszewski JM, Vazquez M, Verschoor S,et al: Rad21-cohesin haploinsufficiency impedes DNA repair and enhances gastrointestinal radiosensitivity in mice. PLoS One 2010, 5:e12112.

24. Moore JM, Rabaia NA, Smith LE, Fagerlie S, Gurley K, Loukinov D, Disteche CM, Collins SJ, Kemp CJ, Lobanenkov VV, Filippova GN: Loss of maternal CTCF is associated with peri-implantation lethality of Ctcf null embryos. PLoS One 2012, 7:e34915.

25. Nora EP, Goloborodko A, Valton AL, Gibcus JH, Uebersohn A, Abdennur N, Dekker J, Mirny LA, Bruneau BG: Targeted Degradation of CTCF Decouples Local Insulation of Chromosome Domains from Genomic Compartmentalization. Cell 2017, 169:930–944 e922.

26. Rao SSP, Huang SC, Glenn St Hilaire B, Engreitz JM, Perez EM, Kieffer-Kwon KR, Sanborn AL, Johnstone SE, Bascom GD, Bochkov ID,et al: Cohesin Loss Eliminates All Loop Domains. Cell 2017, 171:305–320 e324.

27. Schwarzer W, Abdennur N, Goloborodko A, Pekowska A, Fudenberg G, Loe-Mie Y, Fonseca NA, Huber W, C HH, Mirny L, Spitz F: Two independent modes of chromatin organization revealed by cohesin removal. Nature 2017, 551:51–56.

28. Ji X, Dadon DB, Powell BE, Fan ZP, Borges-Rivera D, Shachar S, Weintraub AS, Hnisz D, Pegoraro G, Lee TI,et al: 3D Chromosome Regulatory Landscape of Human Pluripotent Cells. Cell Stem Cell 2016, 18:262–275.

29. Katainen R, Dave K, Pitkanen E, Palin K, Kivioja T, Valimaki N, Gylfe AE, Ristolainen H, Hanninen UA, Cajuso T,et al: CTCF/cohesin-binding sites are frequently mutated in cancer. Nat Genet 2015, 47:818–821.

30. Fujimoto A, Furuta M, Totoki Y, Tsunoda T, Kato M, Shiraishi Y, Tanaka H, Taniguchi H, Kawakami Y, Ueno M,et al: Whole-genome mutational landscape and characterization of noncoding and structural mutations in liver cancer. Nat Genet 2016, 48:500–509.

31. Oti M, Falck J, Huynen MA, Zhou H: CTCF-mediated chromatin loops enclose inducible gene regulatory domains. BMC Genomics 2016, 17:252.

32. Handoko L, Xu H, Li G, Ngan CY, Chew E, Schnapp M, Lee CW, Ye C, Ping JL, Mulawadi F,et al: CTCF-mediated functional chromatin interactome in pluripotent cells. Nat Genet 2011, 43:630–638.

33. Fullwood MJ, Liu MH, Pan YF, Liu J, Xu H, Mohamed YB, Orlov YL, Velkov S, Ho A, Mei PH,et al: An oestrogen-receptor-alpha-bound human chromatin interactome. Nature 2009, 462:58–64.

34. Jin F, Li Y, Dixon JR, Selvaraj S, Ye Z, Lee AY, Yen CA, Schmitt AD, Espinoza CA, Ren B: A high-resolution map of the three-dimensional chromatin interactome in human cells. Nature 2013, 503:290–294.

35. Hnisz D, Day DS, Young RA: Insulated Neighborhoods: Structural and Functional Units of Mammalian Gene Control. Cell 2016, 167:1188–1200.

36. Weinreb C, Raphael BJ: Identification of hierarchical chromatin domains. Bioinformatics 2016, 32:1601–1609.

37. Sofueva S, Yaffe E, Chan WC, Georgopoulou D, Vietri Rudan M, Mira-Bontenbal H, Pollard SM, Schroth GP, Tanay A, Hadjur S: Cohesin-mediated interactions organize chromosomal domain architecture. EMBO J 2013, 32:3119–3129.

38. Tang Z, Luo OJ, Li X, Zheng M, Zhu JJ, Szalaj P, Trzaskoma P, Magalska A, Wlodarczyk J, Ruszczycki B,et al: CTCF-Mediated Human 3D Genome Architecture Reveals Chromatin Topology for Transcription. Cell 2015, 163:1611–1627.

39. Rowley MJ, Nichols MH, Lyu X, Ando-Kuri M, Rivera ISM, Hermetz K, Wang P, Ruan Y, Corces VG: Evolutionarily Conserved Principles Predict 3D Chromatin Organization. Mol Cell 2017, 67:837–852 e837.

40. Stevens TJ, Lando D, Basu S, Atkinson LP, Cao Y, Lee SF, Leeb M, Wohlfahrt KJ, Boucher W, O’shaughnessy-Kirwan A,et al: 3D structures of individual mammalian genomes studied by single-cell Hi-C. Nature 2017, 544:59–64.

41. Seitan VC, Faure AJ, Zhan Y, McCord RP, Lajoie BR, Ing-Simmons E, Lenhard B, Giorgetti L, Heard E, Fisher AG,et al: Cohesin-based chromatin interactions enable regulated gene expression within preexisting architectural compartments. Genome Res 2013, 23:2066–2077.

42. Shen Y, Yue F, McCleary DF, Ye Z, Edsall L, Kuan S, Wagner U, Dixon J, Lee L, Lobanenkov VV, Ren B: A map of the cis-regulatory sequences in the mouse genome. Nature 2012, 488:116–120.

43. Blanchette M, Bataille AR, Chen X, Poitras C, Laganiere J, Lefebvre C, Deblois G, Giguere V, Ferretti V, Bergeron D,et al: Genome-wide computational prediction of transcriptional regulatory modules reveals new insights into human gene expression. Genome Res 2006, 16:656–668.

44. Kim THB, L. O.; Zheng, M; Qu, C; Singer, M. A.; Richmond, T.A.; Wu, Y; Green, R.D.; Ren, B: A high-resolution map of active promoters in the human genome. Nature 2005, 436:876–880.

45. Yanai I, Benjamin H, Shmoish M, Chalifa-Caspi V, Shklar M, Ophir R, Bar-Even A, Horn-Saban S, Safran M, Domany E,et al: Genome-wide midrange transcription profiles reveal expression level relationships in human tissue specification. Bioinformatics 2005, 21:650–659.

46. Kryuchkova-Mostacci N, Robinson-Rechavi M: A benchmark of gene expression tissue-specificity metrics. Brief Bioinform 2017, 18:205–214.

47. Busslinger GA, Stocsits RR, van der Lelij P, Axelsson E, Tedeschi A, Galjart N, Peters JM: Cohesin is positioned in mammalian genomes by transcription,CTCF and Wapl. Nature 2017, 544:503–507.

48. Davidson IF, Goetz D, Zaczek MP, Molodtsov MI, Huis In ‘t Veld PJ, Weissmann F, Litos G, Cisneros DA, Ocampo-Hafalla M, Ladurner R,et al: Rapid movement and transcriptional relocalization of human cohesin on DNA. EMBO J 2016, 35:2671–2685.

49. Nakahashi H, Kwon KR, Resch W, Vian L, Dose M, Stavreva D, Hakim O, Pruett N, Nelson S, Yamane A,et al: A genome-wide map of CTCF multivalency redefines the CTCF code. Cell Rep 2013, 3:1678–1689.

50. Mumbach MR, Rubin AJ, Flynn RA, Dai C, Khavari PA, Greenleaf WJ, Chang HY: HiChIP: efficient and sensitive analysis of protein-directed genome architecture. Nat Methods 2016, 13:919–922.

51. Fuglede B, Topsoe F: Jensen-Shannon Divergence and Hilbert space embedding. IEEE International Symposium on Information Theory 2004, 31.

52. Heidari N, Phanstiel DH, He C, Grubert F, Jahanbani F, Kasowski M, Zhang MQ, Snyder MP: Genome-wide map of regulatory interactions in the human genome. Genome Res 2014, 24:1905–1917.

53. Sugathan A, Waxman DJ: Genome-Wide Analysis of Chromatin States Reveals Distinct Mechanisms of Sex-Dependent Gene Regulation in Male and Female Mouse Liver. Molecular and Cellular Biology 2013, 33:3594–3610.

54. Ling G, Sugathan A, Mazor T, Fraenkel E, Waxman DJ: Unbiased, genome-wide in vivo mapping of transcriptional regulatory elements reveals sex differences in chromatin structure associated with sex-specific liver gene expression. Mol Cell Biol 2010, 30:5531–5544.

55. Wang Z, Zang C, Rosenfeld JA, Schones DE, Barski A, Cuddapah S, Cui K, Roh TY, Peng W, Zhang MQ, Zhao K: Combinatorial patterns of histone acetylations and methylations in the human genome. Nat Genet 2008, 40:897–903.

56. Hay D, Hughes JR, Babbs C, Davies JOJ, Graham BJ, Hanssen L, Kassouf MT, Marieke Oudelaar AM, Sharpe JA, Suciu MC,et al: Genetic dissection of the alpha-globin super-enhancer in vivo. Nat Genet 2016, 48:895–903.

57. Whyte WA, Orlando DA, Hnisz D, Abraham BJ, Lin CY, Kagey MH, Rahl PB, Lee TI, Young RA: Master transcription factors and mediator establish super-enhancers at key cell identity genes. Cell 2013, 153:307–319.

58. Fang B, Everett LJ, Jager J, Briggs E, Armour SM, Feng D, Roy A, Gerhart-Hines Z, Sun Z, Lazar MA: Circadian enhancers coordinate multiple phases of rhythmic gene transcription in vivo. Cell 2014, 159:1140–1152.

59. Iborra FJ, Pombo A, Jackson DA, Cook PR: Active RNA polymerases are localized within discrete transcription ‘factories’ in human nuclei. Journal of Cell Science 1996, 109:1427–1436.

60. Willi M, Yoo KH, Reinisch F, Kuhns TM, Lee HK, Wang C, Hennighausen L: Facultative CTCF sites moderate mammary super-enhancer activity and regulate juxtaposed gene in nonmammary cells. Nat Commun 2017, 8:16069.

61. Hanssen LLP, Kassouf MT, Oudelaar AM, Biggs D, Preece C, Downes DJ, Gosden M, Sharpe JA, Sloane-Stanley JA, Hughes JR,et al: Tissue-specific CTCF-cohesin-mediated chromatin architecture delimits enhancer interactions and function in vivo. Nat Cell Biol 2017, 19:952–961.

62. Osborne CS, Chakalova L, Brown KE, Carter D, Horton A, Debrand E, Goyenechea B, Mitchell JA, Lopes S, Reik W, Fraser P: Active genes dynamically colocalize to shared sites of ongoing transcription. Nat Genet 2004, 36:1065–1071.

63. Martin P, McGovern A, Orozco G, Duffus K, Yarwood A, Schoenfelder S, Cooper NJ, Barton A, Wallace C, Fraser P,et al: Capture Hi-C reveals novel candidate genes and complex long-range interactions with related autoimmune risk loci. Nat Commun 2015, 6:10069.

64. Kung JT, Kesner B, An JY, Ahn JY, Cifuentes-Rojas C, Colognori D, Jeon Y, Szanto A, del Rosario BC, Pinter SF,et al: Locus-specific targeting to the X chromosome revealed by the RNA interactome of CTCF. Mol Cell 2015, 57:361–375.

65. Park HJ, Li J, Hannah R, Biddie S, Leal-Cervantes AI, Kirschner K, Flores Santa Cruz D, Sexl V, Gottgens B, Green AR: Cytokine-induced megakaryocytic differentiation is regulated by genome-wide loss of a uSTAT transcriptional program. EMBO J 2016, 35:580–594.

66. Shukla S, Kavak E, Gregory M, Imashimizu M, Shutinoski B, Kashlev M, Oberdoerffer P, Sandberg R, Oberdoerffer S: CTCF-promoted RNA polymerase II pausing links DNA methylation to splicing. Nature 2011, 479:74–79.

67. Nagano T, Lubling Y, Stevens TJ, Schoenfelder S, Yaffe E, Dean W, Laue ED, Tanay A, Fraser P: Single-cell Hi-C reveals cell-to-cell variability in chromosome structure. Nature 2013, 502:59–64.

68. Wang S, Su JH, Beliveau BJ, Bintu B, Moffitt JR, Wu CT, Zhuang X: Spatial organization of chromatin domains and compartments in single chromosomes. Science 2016, 353:598–602.

69. Vanhille L, Griffon A, Maqbool MA, Zacarias-Cabeza J, Dao LT, Fernandez N, Ballester B, Andrau JC, Spicuglia S: High-throughput and quantitative assessment of enhancer activity in mammals by CapStarr-seq. Nat Commun 2015, 6:6905.

70. Lupianez DG, Kraft K, Heinrich V, Krawitz P, Brancati F, Klopocki E, Horn D, Kayserili H, Opitz JM, Laxova R,et al: Disruptions of topological chromatin domains cause pathogenic rewiring of gene-enhancer interactions. Cell 2015, 161:1012–1025.

71. Raviram R, Rocha PP, Muller CL, Miraldi ER, Badri S, Fu Y, Swanzey E, Proudhon C, Snetkova V, Bonneau R, Skok JA: 4C-ker: A Method to Reproducibly Identify Genome-Wide Interactions Captured by 4C-Seq Experiments. PLoS Comput Biol 2016, 12:e1004780.

72. van de Werken HJ, Landan G, Holwerda SJ, Hoichman M, Klous P, Chachik R, Splinter E, Valdes-Quezada C, Oz Y, Bouwman BA, et al: Robust 4C-seq data analysis to screen for regulatory DNA interactions. Nat Methods 2012, 9:969–972.

73. Thongjuea S, Stadhouders R, Grosveld FG, Soler E, Lenhard B: r3Cseq: an R/Bioconductor package for the discovery of long-range genomic interactions from chromosome conformation capture and next-generation sequencing data. Nucleic Acids Res 2013,41:e132.

74. Melia T, Hao P, Yilmaz F, Waxman DJ: Hepatic Long Intergenic Noncoding RNAs: High Promoter Conservation and Dynamic,Sex-Dependent Transcriptional Regulation by Growth Hormone. Mol Cell Biol 2016, 36:50–69.

75. Servant N, Varoquaux N, Lajoie BR, Viara E, Chen CJ, Vert JP, Heard E, Dekker J, Barillot E: HiC-Pro: an optimized and flexible pipeline for Hi-C data processing. Genome Biol 2015, 16:259.

76. Remeseiro S, Cuadrado A, Carretero M, Martinez P, Drosopoulos WC, Canamero M, Schildkraut CL, Blasco MA, Losada A: Cohesin-SA1 deficiency drives aneuploidy and tumourigenesis in mice due to impaired replication of telomeres. EMBO J 2012, 31:2076–2089.

